# LRRK2 exonic variants associated with Parkinson’s disease augment phosphorylation levels for LRRK2-Ser1292 and Rab10-Thr73

**DOI:** 10.1101/447946

**Authors:** Kenneth V. Christensen, Morten Hentzer, Felix S. Oppermann, Sarah Elschenbroich, Pamela Dossang, Kenneth Thirstrup, Jan Egebjerg, Douglas S. Williamson, Garrick P. Smith

## Abstract

Leucine-rich repeat kinase 2 (LRRK2) is associated to Parkinson’s disease (PD). The most common form of LRRK2 PD is caused by the G2019S variant. Besides G2019S, eight other LRRK2 variants causing familial PD also have amino acid substitutions located in a LRRK2 enzymatic domainsuggesting that enzymatic activity is at the core of mechanisms underlying disease risk. Common LRRK2 polymorphic risk variations such as G2385R, A419V, R1628 and M1646T all reside in other LRRK2 domains. Prior knowledge is limited on how these variants influence LRRK2 function. To investigate the impact on enzymatic function of both rare and common LRRK2 variation a comprehensive profiling of nineteen LRRK2 exonic variants was pursued. Six LRRK2 phosphorylation sites were identified by mass spectrometry. Besides already known phosphorylation sites such as Ser910, Ser935, Ser955, Ser973 and Ser1292 also Thr826 was confirmed by a targeted MRM assay as a LRRK2 phosphorylation site in mammalian cells. Phosphorylation site occupancy for all six LRRK2 sites was obtained but no obvious correlation to risk of disease was found. Instead, application of phospho-specific antibodies targeting LRRK2 phosphorylation sites confirmed that autophosphorylation at Ser1292 was significantly increased for all disease-causing variants whereas no significant differences could be observed for the common intermediate risk variants. Recently, Rab10 and Rab12 have been shown to be bona fide LRRK2 substrates and we find that both rare and common LRRK2 exonic variants augment the phosphorylation of Rab10. This was not observed with Rab12. Furthermore, the protective variant N551K has reduced Rab10 phosphorylation compared to LRRK2 WT. This was not observed with the protective variant R1398H. Our findings support the hypothesis that increased LRRK2 kinase function is associated with increased PD risk but also highlights the need for more sensitive tools for detection of increases in kinase activity in carriers of LRRK2 PD risk variants.

**Abbreviations:**

PD: Parkinson’s disease
LRRK2: leucine-rich repeat kinase 2
MRM: multiple mass spectrometry
MS: mass spectrometry
LC-MS: liquid chromatography mass spectrometry
LOD: limit of detection
MAF: minor allele frequency
CV%: coefficient of variation
SDS-PAGE: SDS-polyacrylamide gel electrophoresis
Roc: Ras of complex
COR: C-terminal of Roc
PRL: pleomorphic risk loci.

## Introduction

Rare genetic variation in the leucine-rich repeat kinase 2 (LRRK2) gene is associated with increased risk of Parkinson’s Disease (PD). To date at least nine rare isoforms of LRRK2 (N1437H, R1441C/H/G/S, Y1699C, S1761R, I2020T and G2019S) causing autosomal dominant familial PD have been identified (1-12). All isoforms have a single non-synonymous amino acid substitution in regions of the LRRK2 gene that encode the core LRRK2 enzymatic domains. This suggests that a perturbed LRRK2 enzymatic function might be involved in pathological mechanisms underlying disease etiology in familial PD. In support of this hypothesis, individuals affected by the most common form of autosomal dominant LRRK2-PD are heterozygous carriers of the G2019S variant encoding a protein with increased kinase activity (13). Further, several LRRK2 disease causing variants including G2019S have shown to increase phosphorylation levels at the LRRK2 autophosphorylation Ser1292 site (14;15). Besides the rare genetic variants several recent reports have associated common variation in the LRRK2 gene both with increased and decreased risk of developing sporadic PD (16-19). In Asian populations the common LRRK2 genomic variants A419V, R1628P and G2385R are associated with increased risk of PD (20-25) whereas in Caucasians the common LRRK2 variant M1646T increases risk of PD (26). Of particular interest, a common haplotype variant that associates with reduced risk of PD has been identified in both Caucasian and Asian populations (26;27). On the protein level the common haplotype contains two non-synonymous substitutions (N551K, R1398H) and one synonymous substitution (K1423K) when compared to LRRK2 WT. All three single nucleotide polymorphisms (SNP) are in strong linkage disequilibrium and therefore co-segregate with the same minor allele frequencies in both populations. Altogether this implies that the enzymatic function of LRRK2 plays a significant role in mechanisms leading to late-onset PD and as such represents a valid target for therapeutic intervention. Current strategies for targeting LRRK2 as a disease-modifying treatment principle in late-onset PD are mostly focused on developing small molecule ATP-competitive inhibitors against the kinase domain (28;29). Currently, there exist no validated animal model of Parkinson’s disease that can be used to predict the clinical efficacy of a LRRK2 inhibitor. LRRK2 G2019S rodent models do not present with Parkinson’s disease symptomatology and although dopaminergic cell death is observed in some models the effects are not fully consistent (30-40). Other approaches used injections of AAV alpha-synuclein or alpha-synuclein seeds in the basal ganglia of rodent LRRK2 transgenic models (41-43). Although the results bare promise a replication study in the AAV alpha-synuclein non-human primate model using a potent and selective LRRK2 inhibitor is warranted. Target populations for LRRK2 therapy are likely to consist of PD patients carrying LRRK2 mutations giving rise to increased LRRK2 kinase activity such as the G2019S populations or by stratifying sporadic PD patients based on LRRK2-kinase activity dependent biomarkers or correlates of such. The most straightforward way would be to stratify patients based on correlates of LRRK2 exonic variation, risk of LRRK2-PD and the level of LRRK2 enzymatic activity. To identify such LRRK2 kinase activity dependent correlates of disease risk we performed a comprehensive analysis of PD-associated LRRK2 exonic variants.

Collectively, the results from proteomic and cell-based studies suggest that even though augmented LRRK2 enzymatic function correlates with the risk of Parkinson’s disease when explored in heterologous expression systems more sensitive detection tools would be a prerequisite when measuring LRRK kinase activity in cellular systems with lower endogenous LRRK2 expression.

## Results

Since the initial cloning of LRRK2 several rare and common LRRK2 exonic variants significantly associated with risk for PD have been identified (Fig 1). Many of these variations are located in the core enzymatic region of LRRK2. Hypothesizing that LRRK2 enzymatic activity is associated with risk of late-onset PD we intended to expand this concept beyond the kinase overactive G2019S variant by carrying out a more comprehensive profiling of PD-associated LRRK2 exonic variants. Besides extensive profiling of the LRRK2 exonic variants in relevant LRRK2 activity-dependent cell-based assays we also aimed at identifying novel LRRK2 autophosphorylation sites, novel LRRK2 interaction partners as well as profiling the impact of both rare and common LRRK2 exonic variation on recently identified LRRK2 kinase activity-dependent substrates such as Rab10 and Rab12 (44). Besides the most compelling disease-linked LRRK2 exonic variants shown in the pleomorphic risk loci map in Fig 1 a number of additional LRRK2 isoforms were also taken into consideration (Table 1), e.g. Y2189C, N2081D and S1647T where ambiguous data on link to disease is reported in the literature (26;45-48). The following non-naturally occurring variants were included as controls in the analyses: the kinase dead mutant D1994A (49;50), the GTPase dead mutant T1348N (51) and the inhibition resistant mutant A2016T (52).

**Figure 1.**
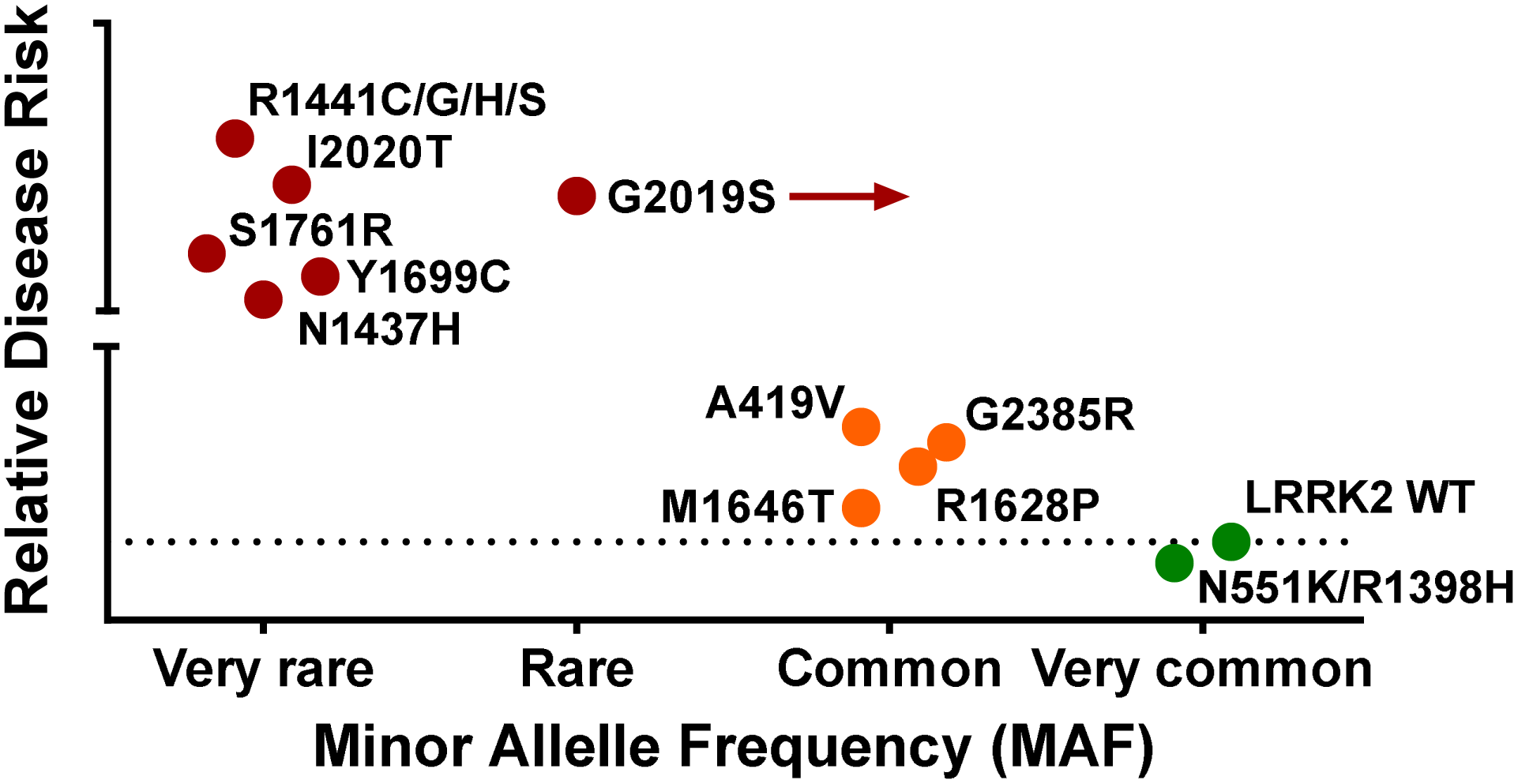
Parkinson’s disease pleomorphic risk loci (PRL) map of LRRK2 exonic variation. The PRL map schematically shows the relative disease risk(s) associated with rare disease-causing LRRK2 variants (red), common LRRK2 variants associated with increased odds-ratios for PD (orange), a common protective haplotype associated with decreased risk of PD (green), and the most frequent LRRK2 variant with a population odds-ratio of 1 (green). The arrow from G2019S indicates that this variant is common in certain ethnic populations such as the Northafrican Berbers or Jews of Ashkenazi decent. The relative odds-ratio of the individual risk variants is based on the publications listed in Table 1. Allele frequencies have been categorized based on data from Ross et al (26). The PRL map for LRRK2 genetic variation was generated with inspiration from Singleton and Hardy, 2011 (78).

**Table 1.**
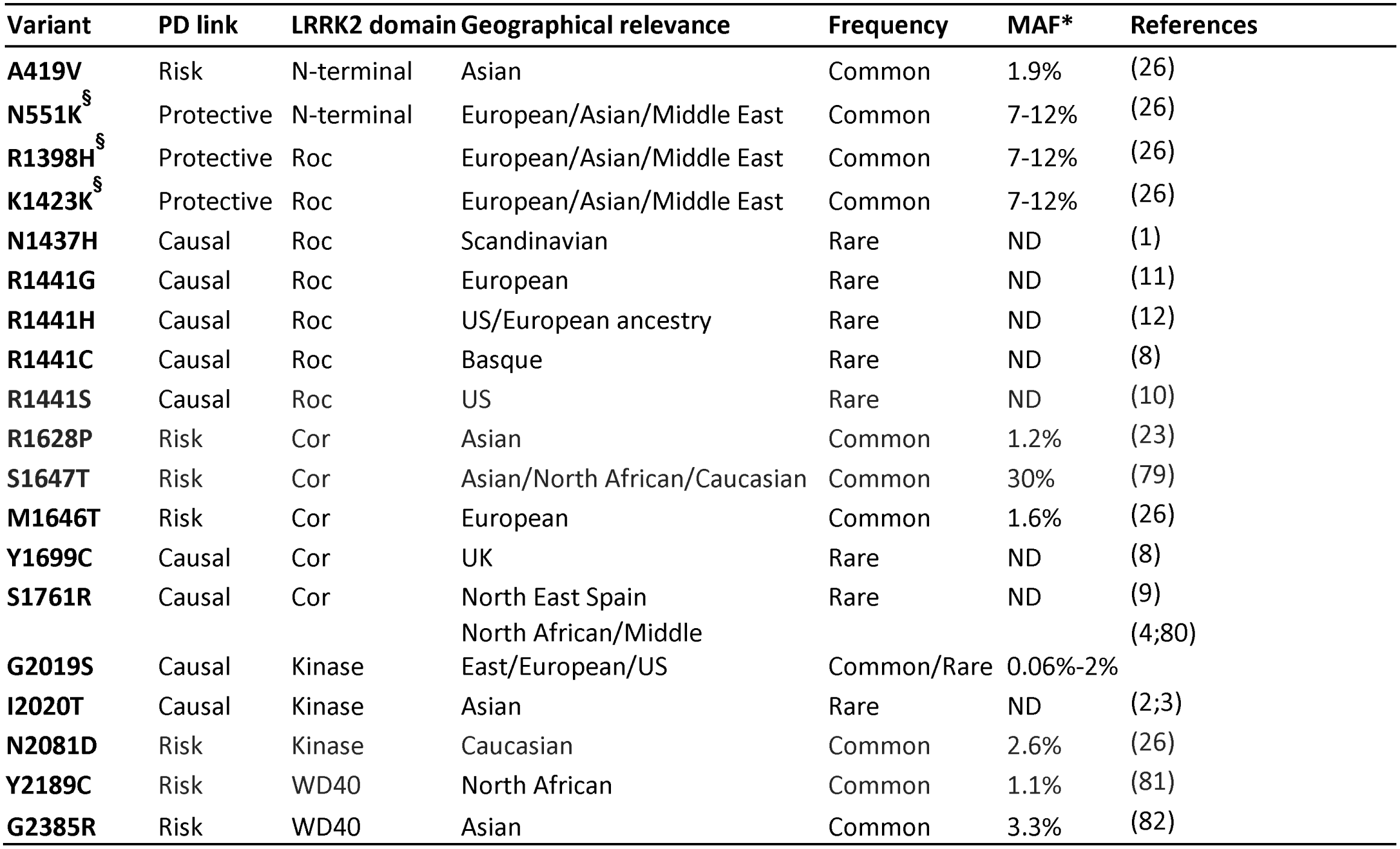
LRRK2 exonic variation and risk of PD.

### Purification of LRRK2 exonic variants

Expression and purification of full-length LRRK2 from bacteria and yeast has so far not proven successful. To identify additional LRRK2 phosphorylation sites cell homogenate from HEK293 transiently transfected with LRRK2 was initially pursued as a LRRK2 protein source. Unfortunately, experiments did not provide sufficient LRRK2 sequence coverage to allow for full stoichiometric analysis of LRRK2 phosphorylation sites by mass spectrometry (MS) (data not shown). Instead, a MS pre-experiment was initiated to determine LRRK2 sequence coverage using mammalian cell purified human LRRK2-G2019S protein. The purified protein contained a small amount of Triton-X100 that potentially could interfere with MS procedures thus two different digest approaches were tested (in-gel or in-solution digest). Both approaches gave good sequence coverage (Table 2); however, the increase in the total number of identified LRRK2 peptides that was observed using the in-solution protocol was largely due to an increased number of miscleaved tryptic peptides. The obtained sequence coverage for LRRK2 upon in-gel digest of 75% was close to the theoretically maximal sequence coverage and met the requirements for subsequent experiments to determine phosphorylation site occupancies. In the subsequent analysis, several phosphopeptides were identified with high confidence (Supplementary Table 1). Among these were identified phosphopeptides containing the Ser910, Ser935, Ser955 and Ser973 sites known to be phosphorylated in cells and *in vivo* (53-57). Thus, mammalian cell purified LRRK2 protein proved useful for assessing LRRK2 phosphorylation patterns with good sequence coverage.

**Table 2.** Comparison of in-gel and in-solution digest methods.

| Digest method | Sequence coverage | Peptides | Miscleaved peptides | Miscleaved peptides (%) |
| --- | --- | --- | --- | --- |
| In-gel | 74.9 | 186 | 63 | 33.9 |
| In-solution | 75.5 | 257 | 148 | 57.6 |

### Production and characterization of LRRK2 exonic variants

Consequently, LRRK2 protein purified from mammalian cells of the nineteen (19) LRRK2 exonic variants plus additional batches of G2019S and LRRK2 WT were obtained (see Methods). In total, 22 batches of LRRK2 were investigated. The list of investigated LRRK2 variants comprised two (2) batches of LRRK2 WT, three (3) batches of G2019S, fourteen (14) exonic variants and three (3) additional mutants. The latter three were used to determine the effect of mutating amino acids vital for LRRK2 function in the core enzymatic domains. The concentration, purity and specific activity data as well as batch information for all the LRRK2 exonic variants can be found in Supplementary Table 2.

### Phosphoproteomic analysis of individual variants

Prior to determination of phosphorylation site stoichiometries a comprehensive phosphopeptide reference dataset was established for LRRK2. Protein samples were digested according to the in-gel protocol and 2.5 µg of each variant were utilized for phosphopeptide enrichment and mass spectrometry analysis. The list of identified phosphorylation sites was filtered for LRRK2-derived phosphorylation sites that could be localized with high confidence, i.e. class-I phosphorylation sites, indicated by a localization probability ≥ 0.75 and a score difference ≥ 5 of class-I (58). Altogether, 43 LRRK2-derived phosphorylation sites were identified (Supplementary File 1). The list comprises multiple known phosphorylation sites, e.g. the autophosphorylation site Ser1292 as well as the indirect phosphorylation sites Ser910, Ser935, Ser955 and Ser973. Interestingly, the Thr1410 site was also observed; a phospho-site which has not been previously been confirmed in cell-based assays but only observed after subjecting cell lysates to an auto-phosphorylation assay or by incubating purified LRRK2 protein under cell-free conditions in the presence of saturating amounts of ATP (59-61). To our surprise, phosphorylation at Ser1444, another reported phospho-site, was only picked up for the R1441H variant site (9%). Situated in the ROC domain the Ser1444 site modulates LRRK2 interaction with 14-3-3 proteins and it has previously been confirmed as a target for PKA phosphorylation using ROC single-domain constructs (62).

Estimation of the phosphorylation site occupancy for the individual variants was performed using the in-gel digest protocol. For each variant one aliquot was incubated with alkaline phosphatase whereas the other aliquot was control-incubated without phosphatase (Fig 2). To enable quantitative LC-MS/MS analysis the two samples were then differentially labelled with mTRAQ reagent and combined before LC-MS/MS analysis. The quantified ratios for non-phosphorylated peptides directly correlate with the level of phosphorylation of the respective peptide (Supplementary File 1). Thus, on the phosphopeptide reference dataset it is possible to conclude likely phosphorylation site(s) that are associated to the quantified peptides. A heat map of the phosphorylation occupancies and positions for all LRRK2 exonic variants indicate a cluster of higher phosphorylation in a distinct region of the LRRK2 protein ranging between the ANK- and LRR domains (Fig 3). This region comprises Ser910, Ser935, Ser955 and Ser973 which are known cellular LRRK2 phosphosites that are critical for interaction with 14-3-3 proteins (53;54;63). Indeed, peptides from several 14-3-3 proteins were also found to be significantly enriched in the samples as the in-gel LC-MS/MS approach also allowed for identification of peptides that co-purified with LRRK2. Only exemptions were the GTPase-dead variant T1348N and the causal disease variant N1437H where co-purification of 14-3-3 proteins were substantially diminished compared to LRRK2 WT (Supplementary Fig 1). These findings highlight the importance of the GTPase function for phosphorylation of LRRK2 and concomitant interaction with 14-3-3. In addition, the VEPSWLGPLFPDKTSNLR peptide (amino acid: 813-830) was revealed to be highly phosphorylated across most LRRK2 variants. The peptide includes Thr826 that previously has been identified as an in vitro autophosphorylation site (61).

**Figure 2.**
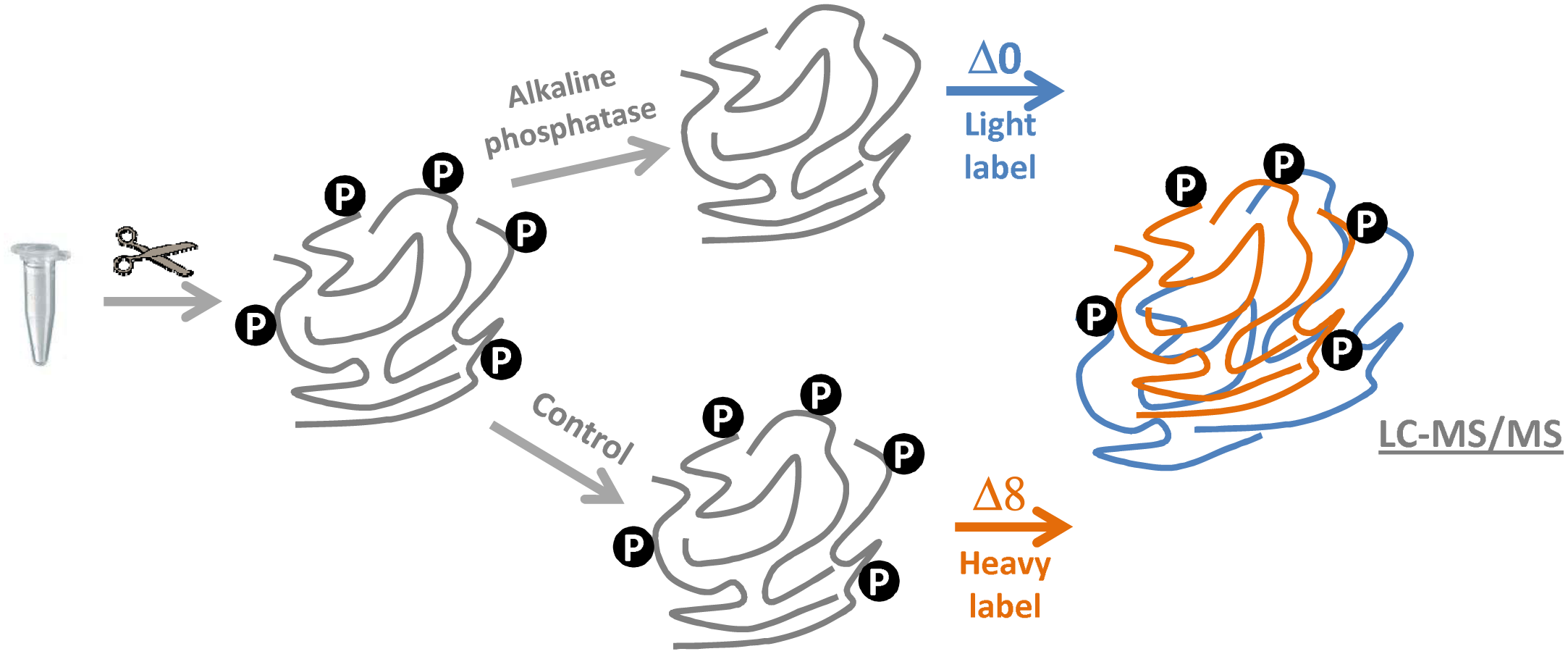
Flowchart of the phosphoproteomic study. LRRK2 protein samples were digested in-gel with trypsin and the desalted peptides were aliquoted. For the enzymatic dephosphorylation of phosphopeptides one aliquot was incubated with alkaline phosphatase (PHOSPHO) whereas the other aliquot was control-incubated without phosphatase (CTRL). To enable quantitative LC-MS/MS analysis the two samples were then differentially labeled with mTRAQ reagent(s) and combined before LC-MS/MS analysis. The quantified ratios for non-phosphorylated peptides directly correlate with the level of phosphorylation of the respective peptide. Phosphorylation stoichiometries were calculated according to the following equation: % phosphorylation = (Light^Δ0^-Heavy^Δ8^)/Light^Δ0^.

**Figure 3.**
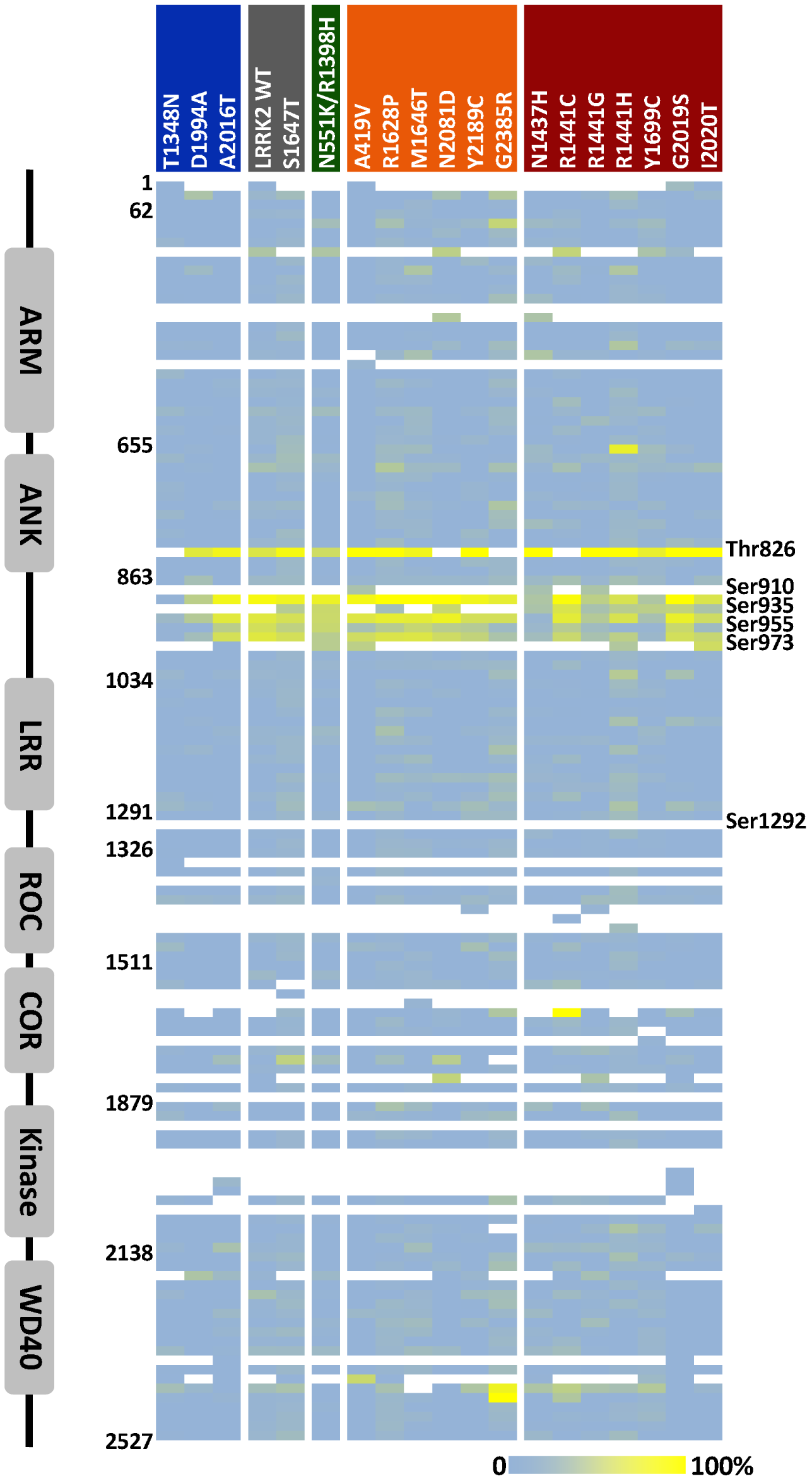
Phospho-site stochiometries of LRRK2 exonic variants. *Right*, heat map of the calculated phosphorylation site occupancy for all LRRK2 peptides across the nineteen LRRK2 exonic variants. The color coding is a gradient from no phosphorylation (0%) in blue to a maximum level of phosphorylation (100%) in yellow. The LRRK2 variants have the following color coding: disease-causing variants (red), common risk variants (orange), protective variant (green), wild type variants (grey) and other variants (blue). *Left,* image of the LRRK2 primary structure with accompanying amino acid positions relative to the structural domains. Adapted from Christensen et al, 2017 (28).

Phosphorylation at the Ser1292 site could not be determined; most likely due to the existence of two tryptic cleavage sites in close proximity to the Ser1292 residue. A one-way ANOVA analysis was applied on all peptides being quantified in at least two out of four replicate experiments for the respective LRRK2 variant. The analysis revealed that the three peptides harboring the Ser910, Ser935 and Ser973 phosphorylation sites, respectively, exhibited a significantly different level of phosphorylation site occupancy for the individual LRRK2 variants. However, due to large variability in the determined phosphorylation site occupancies for these five phosphorylation sites an assessment of intra-individual differences between LRRK2 exonic variants was not possible (see Supplementary Fig 2-3). For that purpose, a more sensitive approach by targeted mass spectrometry is needed.

### Stochiometric analysis of phosphorylation sites by MRM

Next, we developed and applied multiple reaction monitoring mass spectrometry (MRMMS) assays to the purified LRRK2 protein samples. If sensitive enough such MRM assays could be relevant for measuring LRRK2 kinase-activity dependent phosphorylation in relevant human bio samples. Of relevance, would be carriers of G2019S as well as from individuals carrying selected LRRK2 variants associated with risk of PD.

Identified phosphorylation sites with a high degree of phosphorylation occupancy as well as the peptide comprising the Ser1292 site were selected for targeted MRM-based quantification and MRM assays were developed for the corresponding nonphosphorylated peptides (Table 3). For the latter phosphorylation site, alternative enzymatic digest protocols were assessed and it was found that endoproteinase GluC digest yielded a peptide suitable for MS-analysis of Ser1292; both an oxidized and non-oxidized version of the target peptide sequence were included. Further experimental evidence suggested that the non-oxidized peptide was prone to fast oxidization and hence not stable in solution. Thus, quantification was accomplished using the oxidized peptide only. Since many of the peptides harbored more than one putative phosphorylation site the same indirect strategy as for the initial LC-MS/MS experiments was pursued. Thus, MRM assays were developed for the non-phosphorylated peptide and applied onto sample aliquots with or without phosphatase treatment. Assay characteristics for both the tryptic-peptide MRM assays and the GluC-peptide MRM assay were established. All peptides showed acceptable LOD values and assays proved to be linear over the tested concentration range (Table 3 and Supplementary Fig 5-6).

**Table 3.** LRRK2 MRM assay characteristics.

| **LRRK2**<br>**Phosphorylation site(s)** | **LRRK2**<br>**Peptide sequence** | **Enzyme**<br>**Digest** | **Calculated LOD**<br>**(amol/μl)** |
| --- | --- | --- | --- |
| T826 | VEPSWLGPLFPDKTSNLR | Tryptic | 67,87 |
| S908; S910; S912 | SNISISVGEFYR | Tryptic | 52,19 |
| S933; S935 | HSNSLGPIFDHEDLLK | Tryptic | 579,26 |
| S954; S955; S958 | ILSSDDSLR | Tryptic | 29,13 |
| S971; S973; S975; S976; S979 | HSDSISSLASER | Tryptic | 44,94 |
| S1292 | MGKLSKIWDLPLDE | GluC | ND |
| S1292 | M(ox)GKLSKIWDLPLDE | GluC | 29,22 |

To evaluate the usefulness of the MRM-MS assay approach, only a subset of the most relevant LRRK2 exonic variants including G2019S, the common risk-associated LRRK2 exonic variants such as M1646T, A491V, G2385R as well as the protective variant N551K/R1398H were profiled using the MRM assays. LRRK2 WT and the kinase-dead D1994A were also included as controls in the analysis. Protein samples were aliquoted with one aliquot treated with alkaline phosphatase and the other aliquot with buffer only. Subsequently, the amount of non-phosphorylated peptide was quantified for both samples. Using this approach, the MRM assays delivered much more precise data than obtained in the LC-MS/MS study performed with data dependent MS/MS acquisition. Confidence intervals for all investigated peptides were significantly smaller using the MRM approach (Fig 4 and Supplementary File 2). For the four peptides harboring the following putative phosphorylation sites (S908/S910/S912), (S933/S935), (S954/S955/S958), and (S971/S973/S975/S976/S979), the percentage of phosphorylation could be determined with very little variance for all variants except for the kinase-dead variant D1994A. Hence, even small differences in phosphorylation occupancy between these LRRK2 variants could be evaluated. For the phosphorylation sites Thr826 and Ser1292 (the latter obtained by GluC-digest), however, the variance was higher. Most phosphorylation site levels are in the following order G2019S≥LRRK2 WT=M1646T>A419V=N551K/R1398H>>G2385R= D1994A. The only exception is for the Ser955 site where levels of A419V are lower than the protective variant N551K/R1398H.

**Figure 4.**
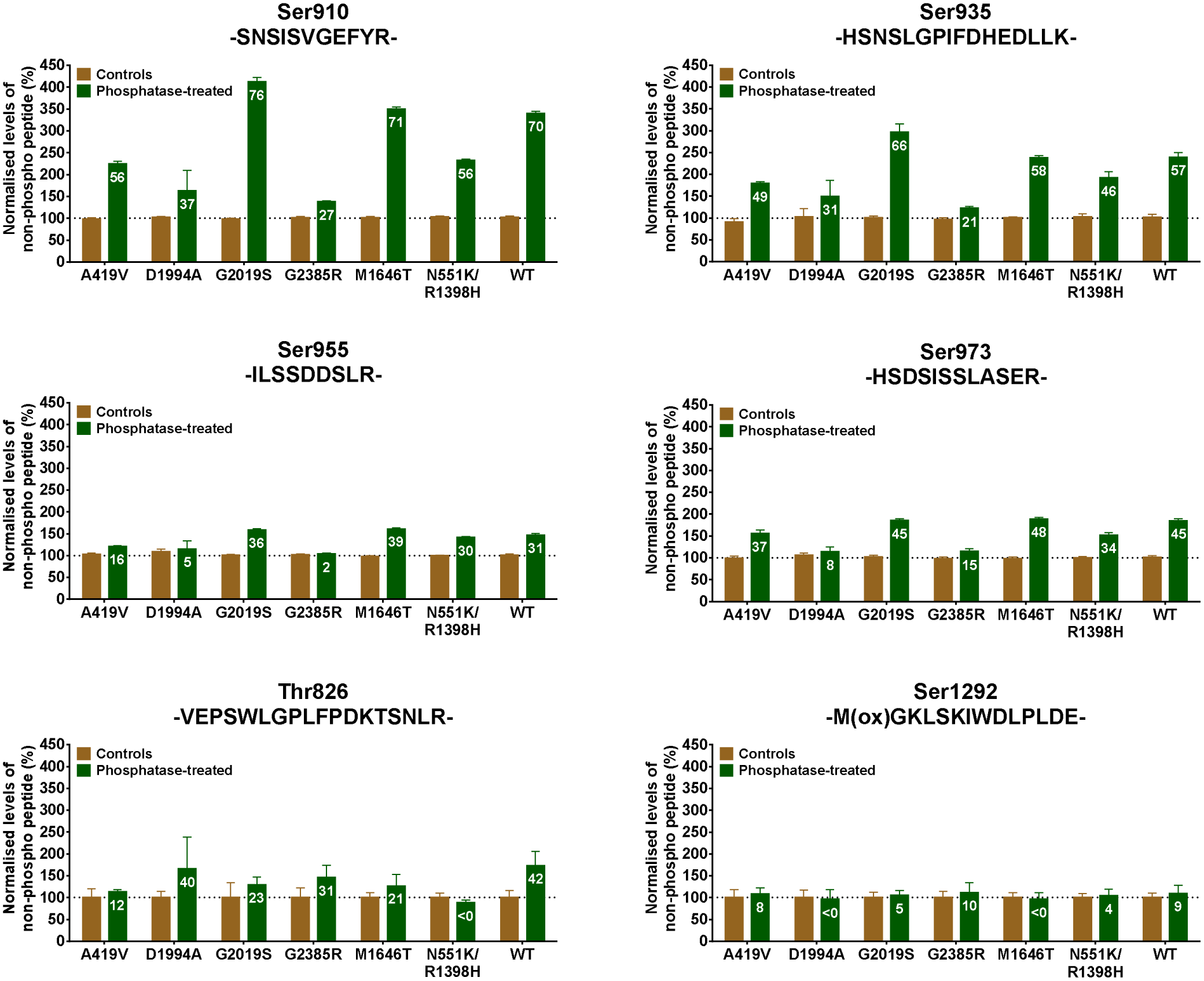
Relative LRRK2 phosphorylation site occupancy (%) determined by MRM. LRRK2 exonic variants were in-gel digested and resuspended in buffer containing heavy-labeled non-phosphorylated peptides from each of the developed six MRM assays (containing Ser910, Ser935, Ser955, Ser973, Ser826 and Ser1292, respectively). For each assay, a sample aliquot was treated with alkaline phosphatase (green) and an aliquot was control incubated with buffer only (brown). After the MS analysis, total area ratios between endogenous peptide and isotopically labeled peptide were used to calculate the phospho-site occupancy for LRRK2 variant. Data is presented as normalized means ± SEM (n=6). The number on the green bars denotes the calculated % phosphorylation of the peptide for the respective LRRK2 variant. The peptide sequence and most likely LRRK2 phospho-site is indicated above the graphs.

The analysis highlights that the order of the phosphorylation site occupancy is as follows: pSer910>pSer935>pSer973>pSer955>Thr826>>pSer1292. For LRRK2 WT the approximate phospho-site occupancies for Ser910, Ser935, Ser955, Ser973 and Ser826 were 70%, 60%, 30%, 45% and 40%, respectively. For these five sites, there are no obvious pattern related to disease risk. The observed phosphorylation at the Ser1292 site is much less than for any of the other sites, and, for most variants, levels are below 10%. Given the variability in MRM-based quantification of the corresponding Ser1292 peptide, detection of significant differences between variants cannot be accomplished at this low stoichiometry. Thus, more sensitive and precise assays such as phospho-specific antibody based analyses are required.

### Western Blot analysis of cellular LRRK2 phosphorylation

LC-MS/MS and MRM-MS studies have confirmed cell-based phosphorylation of six LRRK2 phosphorylation sites at Thr826, Ser910, Ser935, Ser955, Ser973 and Ser1292. However, the MS-based analyses showed large variability in the determination of phosphorylation occupancies not allowing for proper evaluation of the impact of exonic variation on LRRK2 phosphorylation patterns. The phospho-specific antibodies targeting the phosphorylation sites at Ser910, Ser935, Ser955, Ser973 and Ser1292 provide such needed alternatives. Thus, a Western Blot based approach was implemented looking at the difference between LRRK2 exonic variants with respect to phosphorylation. Initially, 10 ng of purified LRRK2 protein was loaded in duplicate on the SDS gels and total LRRK2 and phospho-LRRK2 subsequently quantified by Western Blot. For normalization purposes LRRK2 WT and the kinase-dead variant D1994A was loaded in duplicate or triplicate on each gel (Fig 5A-D). The experiments were repeated three times. For the cellular phosphorylation sites Ser910, Ser935, Ser955 and Ser973 the results were similar to what was observed for the seven (7) variants that underwent MRM-based quantification. The phosphorylation at these four (4) sites seems to be dependent on an ATP- or GTP-bound LRRK2 kinase as both the kinase-dead D1994A and the GTPase-dead variant T1348N have no or very low levels of phosphorylation at this site (Fig 6A-D). All rare disease variants with mutations in the GTPase domain also show a significant lower phosphorylation levels for Ser910/Ser935/Ser955/Ser973 when compared to LRRK2 WT. Although not as prominent an effect the same is observed for many of the common risk variant as well as for the protective variant N551K/R1398H for Ser910/Ser935/Ser955. The kinase-overactive variant G2019S has a similar phosphorylation pattern to LRRK2 WT with respect to Ser935 and Ser973 whereas for Ser910 and Ser955 G2019S has a significant lower phosphorylation. Thus, it appears that for these variants there is no obvious phosphorylation pattern at Ser910/Ser935/Ser955/Ser973 that seems to correlate with the disease risk which is also consistent with literature reports (55).

**Figure 5.**
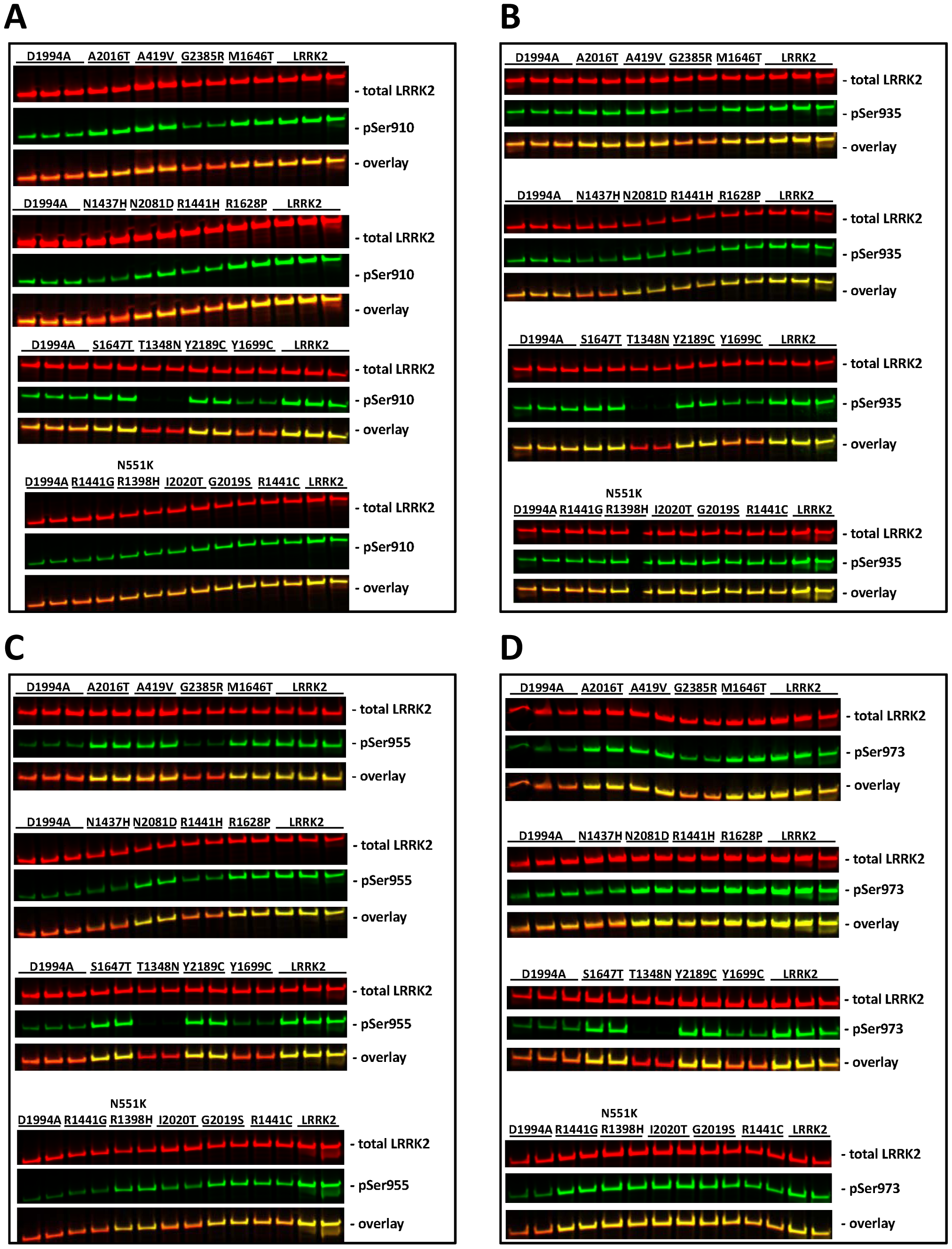
Westen blot determination of LRRK2 phosphorylation at LRRK2 exonic variants. Mammalian purified LRRK2 protein (10ng) from the individual LRRK2 exonic variant samples were probed by SDS-PAGE and subsequent quantitative Western blotting. A-D, LiCor Western Blot image example showing mammalian purified total LRRK2 immunoreactivity (red panels) and A, LRRK2-pSer910; B, LRRK2-pSer935; C, LRRK2-pSer955; D, LRRK2-pSer973 immunoreactivity (green panels) as well as overlay for the various LRRK2 exonic variants. Full-length blots are presented in Supplementary Fig 2.

**Figure 6.**
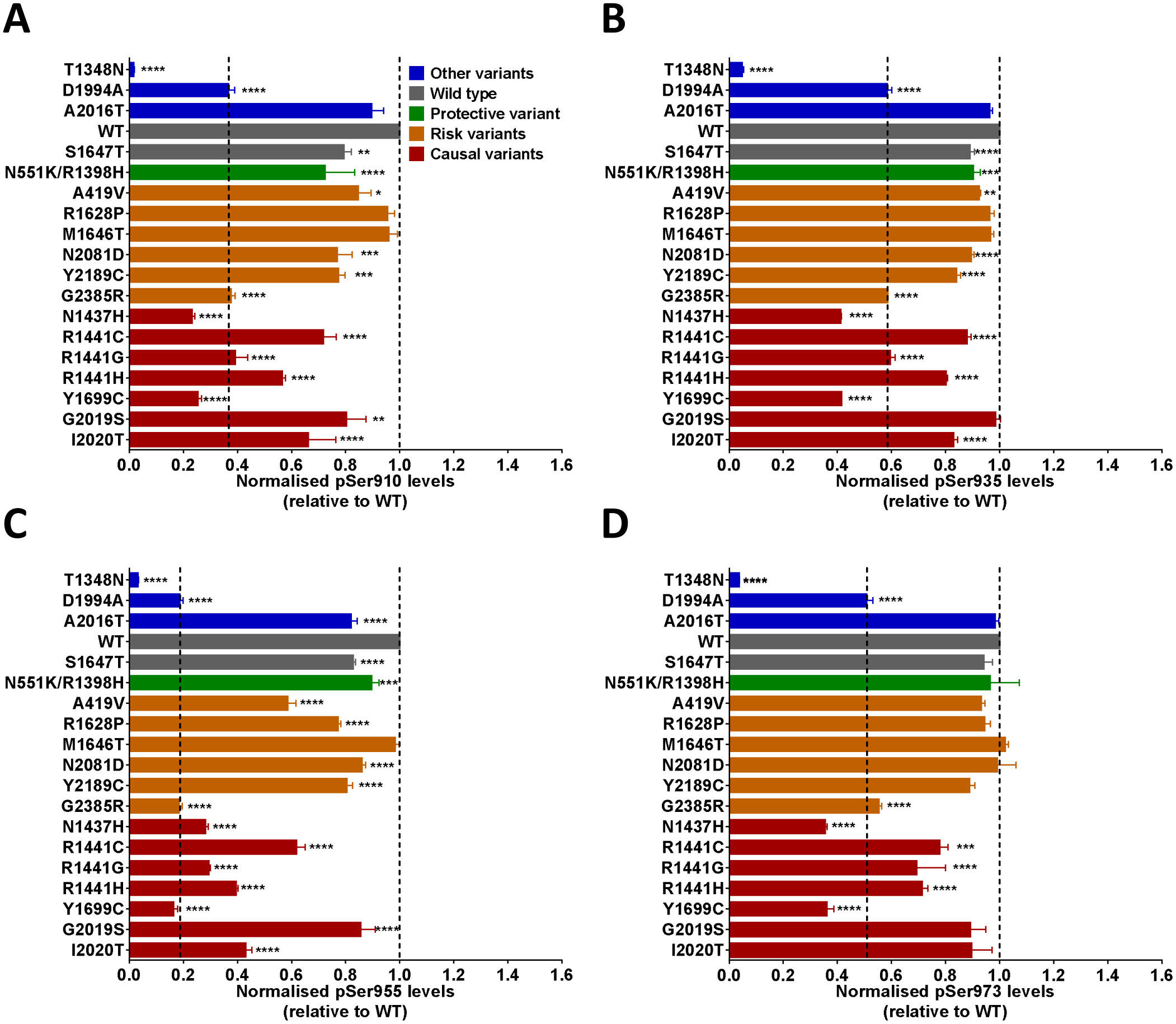
Quantification of LRRK2 exonic variant phosphorylation. A, normalized LRRK2-pSer910/total LRRK2 ratio (n=3 experiments). B, normalized LRRK2-pSer935/total LRRK2 ratio (n=3 experiments). C, normalized LRRK2-pSer955/total LRRK2 ratio (n=3 experiments). D, normalized LRRK2-pSer973/total LRRK2 ratio (n=3 experiments). Data was analyzed by two-way ANOVA with Holm-Sidak’s multiple comparisons test. Data presented as means ± SEM; *p<0.05; **p<0.01; ***p<0.001; ****p<0.0001 vs. WT.

### Western Blot analysis of LRRK2 autophosphorylation at Ser1292

The sensitivity of the Odyssey CLx based pSer1292 assay was close to the detection limit when only loading 10 ng of recombinant LRRK2 protein. Optimization efforts showed that the detection limit of total LRRK2 protein is independent of LRRK2 variant (data not shown). On the contrary, the detection limit for Ser1292 phosphorylation at G2019S is approximately 1ng using Western Blot-based measures whereas for LRRK2 WT at least 30ng recombinant protein is required before increased Ser1292 phosphorylation over background and kinase-dead D1994A levels can be detected (Supplementary Fig 4). SDS-PAGE with 90ng of LRRK2 protein loaded allowed for subsequent Western blot evaluation of the Ser1292 phosphorylation status of the common LRRK2 exonic variants (Fig 7A). Results obtained using these conditions indicate that all disease causing LRRK2 exonic variants have higher relative Ser1292 phosphorylation when compared to LRRK2 WT and the protective variant N551K/R1398H (Fig 7B, red variants). A one-way ANOVA shows a statistically significant difference between group means (p<0.0001; n=3) and a post-hoc Dunnett’s Multiple Comparison test shows that the Ser1292 phosphorylation at LRRK2 WT is significantly different from the Ser1292 phosphorylation at all the rare disease causing LRRK2 variants I2020T, G2019S, N1437H, R1441C, R1441G, R1441H and Y1699C. Thus, autophosphorylation at the Ser1292 site seems to correlate with risk of disease. Additional experimental comparison between the pSer1292 levels at the kinase-dead LRRK2 variant D1994A and LRRK2 WT shows (Fig 7C) that there is a statistically significant difference between the pSer1292 levels at the kinase-dead LRRK2 variant D1994A and LRRK2 WT (paired t-test, p<0.0001; n= 16). This combined with the fact that pSer1292 levels at G2019S is ∼6-fold higher than LRRK2 WT strongly suggests that phosphorylation at the Ser1292 site is both dependent on as well as correlates well with LRRK2 kinase activity. No significant differences between LRRK2 WT and common risk variants were observed.

**Figure 7.**
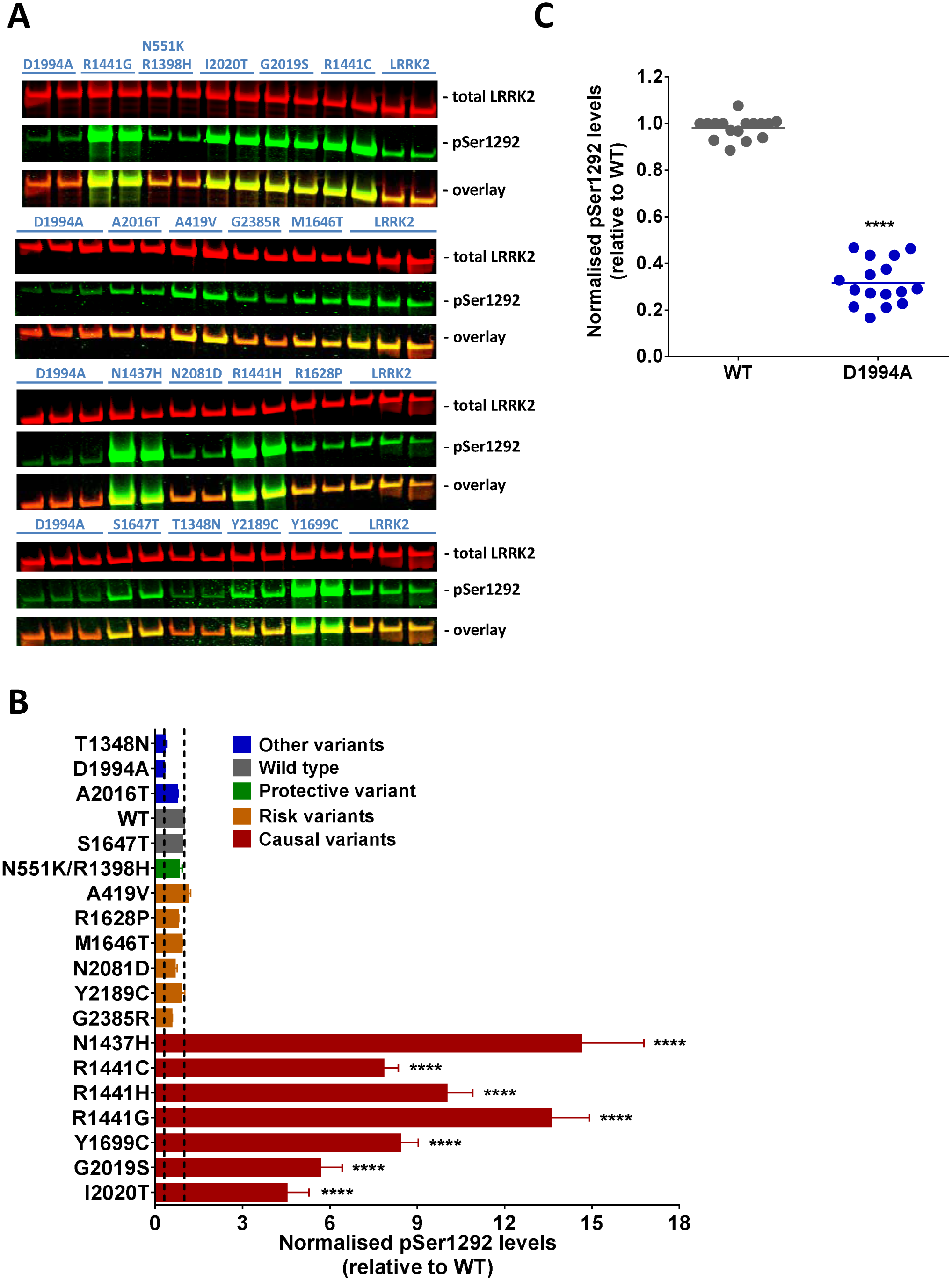
Quantification of LRRK2-Ser1292 phosphorylation in LRRK2 exonic variants. Mammalian purified LRRK2 protein (90ng) from the individual LRRK2 exonic variant samples were probed by SDS-PAGE and subsequent quantitative Western blotting. A, LiCor Western Blot image example showing mammalian purified total LRRK2 immunoreactivity (red panels) and LRRK2-pSer1292 immunoreactivity (green panels) as well as overlay for the various LRRK2 exonic variants. Full-length blots are presented in Supplementary Fig 2. B, the kinase-dead LRRK2 D1994A variant is significant less phosphorylated on Ser1292 when compared to LRRK2 WT. Data presented as means ± SEM; paired t-test (n=16). C, all rare disease-causing LRRK2 exonic variants have increased levels of LRRK2-pSer1292. Data was analyzed by one-way ANOVA with Dunnett’s multiple comparisons test. Data presented as means ± SEM (n=4); *p<0.05; **p<0.01; ***p<0.001; ****p<0.0001 vs. WT.

### Characterization of the Thr826 phosphorylation site

So far development of phospho-specific antibodies targeting Ser910, Ser935, Ser955, Ser973 and Ser1292 have successfully been pursued and commercialized via the Michael J. Fox Foundation (64;65); however, a specific antibody targeting phosphorylation at the Ser826 site is not commercially available. Thus, production of a polyclonal phosphospecific antibody targeting LRRK2 phosphorylated Thr826 was initiated (see Materials and Methods). However, the antibody affinity was not sufficient high to be able to detect phosphorylation at Thr826 in HEK293 cells transiently expressing various LRRK2 exonic variants or by Western Blot-based assessment of mammalian purified LRRK2 exonic variant proteins (data not shown).

### Western Blot analysis of Rab10 and Rab12 phosphorylation

Rab10 and Rab12 were recently identified and validated as human LRRK2 substrates (44). To evaluate whether LRRK2-dependent phosphorylation of Rab10 and Rab12 correlates with risk of disease associated with LRRK2 variation two validated phospho-specific polyclonal antibodies targeted against Rab10 phosphorylated at Thr73 and Rab12 phosphorylated at Ser106, respectively were used to measure Rab10 and Rab12 phosphorylation in lysates from HEK293 transiently co-expressing the respective Rab GTPase as well as the individual LRRK2 exonic variants (66). Subsequent Western Blot based analyses of relative Rab10-Thr73 phosphorylation show that LRRK2 WT phosphorylates HA-tagged Rab10 but not the kinase dead D1994A nor the GTPase dead T1348N variants (Fig 8A-C). Increased levels of phosphorylation are observed in cells expressing the causal disease variants I2020T, R1441C/G/H, Y1699C and G2019S when compared to cells expressing LRRK2 WT; the latter variant, although, to a lesser extent (Fig 8C). Interestingly, when compared to cells expressing LRRK2 WT, and in contrast to the pSer1292 data, increased Rab10-pThr73 levels were observed in cells expressing the common LRRK2 disease risk variants such as R1628P, A419V, G2385R, N2081D and M1646T. In addition, the protective LRRK2 exonic variants N551K and R1398H show opposing effects. N551K has reduced and R1398H has increased pThr73 levels when compared to LRRK2 WT. In the experiments, there was no significant difference in total LRRK2 levels between variants (one-way ANOVA; F (17, 54) = 1.100, p = 0.38). Similarly, total Rab10 levels showed no statistically significant differences between variants (one-way ANOVA; F (17, 54) = 1.057, p = 0.42). For Rab12 the Western Blot-based results show that LRRK2 WT phosphorylates HA-tagged Rab12 but not the kinase dead D1994A nor the GTPase dead T1348N variants (Fig 9A-C). In contrast to the Rab10 data elevated levels of phosphorylation compared to LRRK2 WT were not observed in cells expressing the causal disease variants nor was this observed for the common LRRK2 disease risk variants. Similarly, the protective LRRK2 exonic variants N551K and R1398H showed no decreased or increased Rab12 phosphorylation compared to LRRK2 WT. There was a significant difference in the total LRRK2 levels between variants (one-way ANOVA; F (17, 45) = 2.772, p <0.01). A post-hoc analysis showed significant increased LRRK2 expression in cells expressing N2081D when compared to LRRK2 WT (Holm-Sidak’s multiple comparisons test; p<0.01). There was no significant difference in total Rab12 levels between variants (one-way ANOVA; F (17, 54) = 1.036, p = 0.44). Interestingly, the LRRK2-pSer935 data seems to deviate between Rab10 and Rab12 expressing cells. In both studies, the kinase and GTPase dead variants have a statistically significant decreased Ser935 phosphorylation compared to LRRK2 WT (Fig 8D and 9D). However, in Rab10 expressing cells there is also a statistically significant lower Ser935 phosphorylation in all disease variants whereas pSer935 levels of the protective variants are similar to LRRK2 WT (Fig 8D). In contrast, for Rab12 expressing cells the relative LRRK2-pSer935 levels for all variants are generally more similar to LRRK2 WT (Fig 9D). A couple of variants such as G2385R and R1628P have reduced levels and the LRRK2 protective variants N551K and R1398H have increased levels; however, besides the kinase and GTPase dead variants none of the changes in Rab12 expressing cells are statistically significant different from the pSer935 levels for LRRK2 WT.

**Figure 8.**
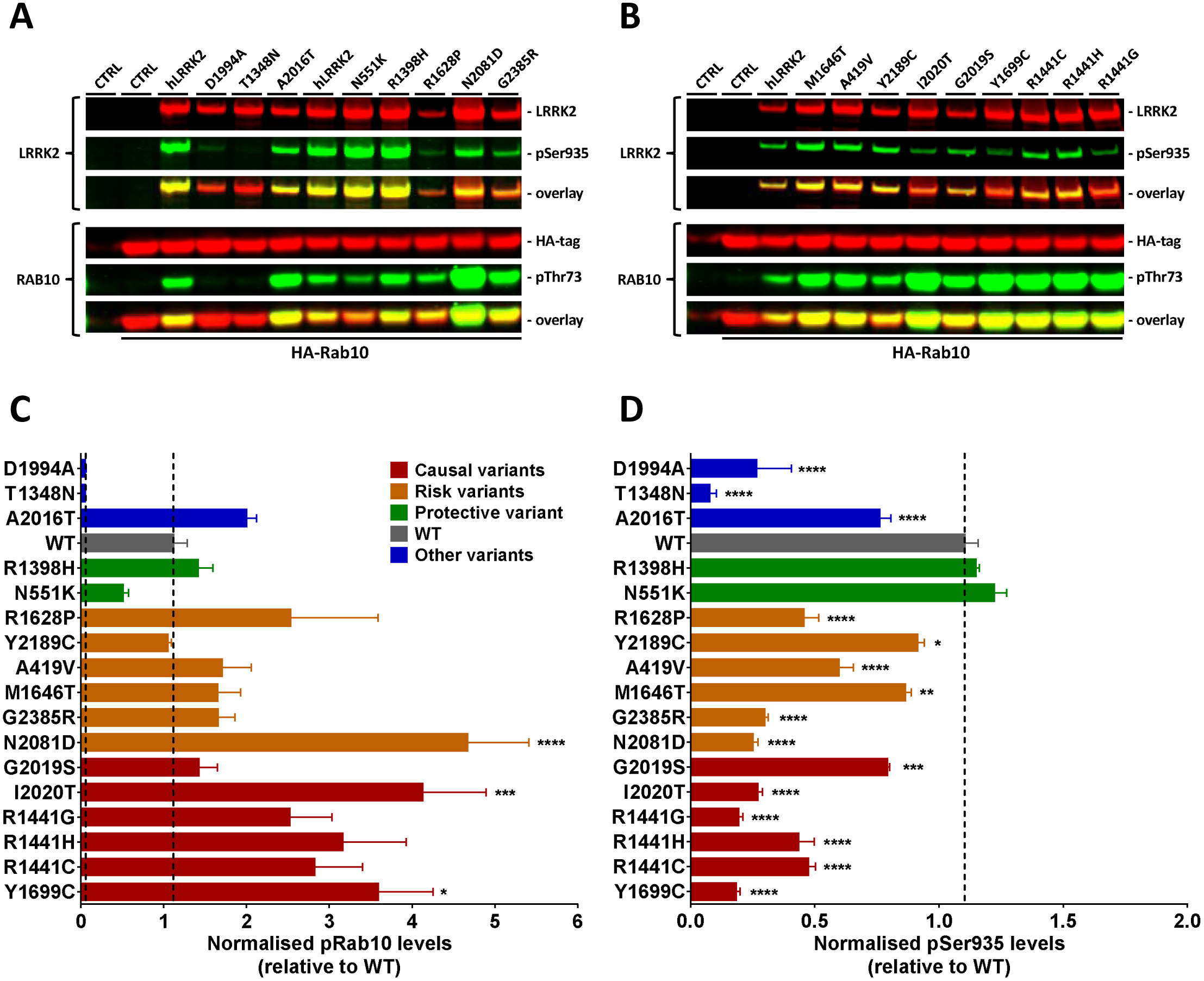
Quantification of LRRK2 kinase-dependent Rab10-Thr73 phosphorylation in HEK293 cells. *A1-2*, LiCor Western Blot image examples showing overexpressed total LRRK2 and HA-tagged Rab10 immunoreactivity (red panels), LRRK2-pSer935 and Rab10-pThr73 immunoreactivity (green panels) as well as overlay in cells overexpressing various LRRK2 exonic variants with HA-tagged wild type Rab10. *Top panels*, LRRK2 and LRRK2-pSer935. *Bottom panels*, Rab10 and Rab10-pThr73. Full-length blots are presented in Supplementary Fig 2. ***B***, quantification of normalized Rab10-pThr73/total Rab10 ratio (n=4 experiments). ***C***, quantification of normalized LRRK2-pSer935/total LRRK2 ratio (n=4 experiments). Data was analyzed by one-way ANOVA with Holm-Sidak’s multiple comparisons test. Data presented as means ± SEM; *p<0.05; **p<0.01; ***p<0.001; ****p<0.0001 vs. LRRK2 WT.

**Figure 9.**
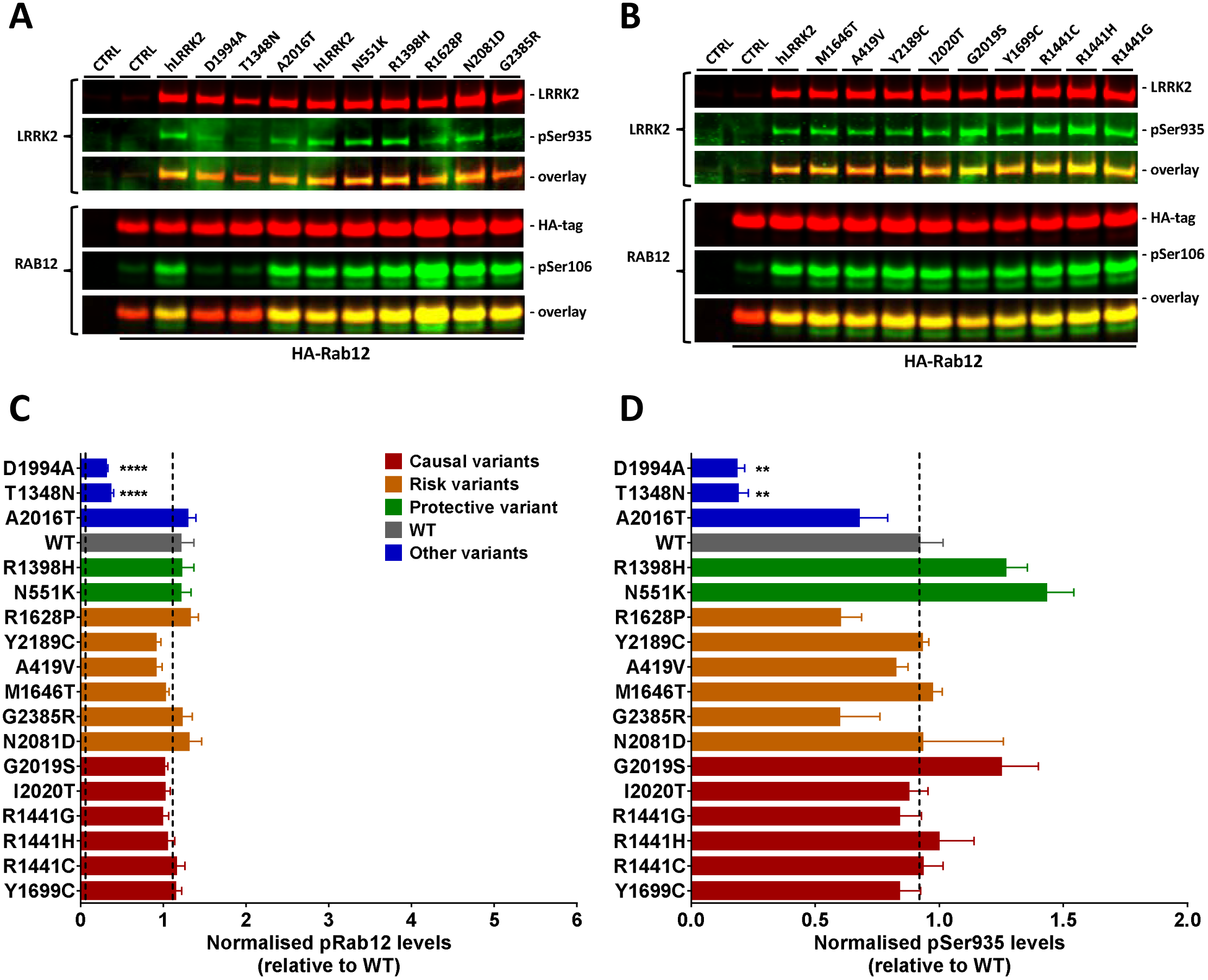
Quantification of LRRK2 kinase-dependent Rab12-Ser106 phosphorylation in HEK293 cells. A1-2, LiCor Western Blot image showing overexpressed total LRRK2 and HA-tagged Rab12 immunoreactivity (red panels), LRRK2-pSer935 and Rab12-pSer106 immunoreactivity (green panels) as well as overlay in cells overexpressing various LRRK2 exonic variants with either HA-tagged wild type Rab12. Top panels, LRRK2 and LRRK2-pSer935. Bottom panels, Rab12 and Rab12-pSer106. Full-length blots are presented in Supplementary Fig 3. B, quantification of normalized Rab12-pSer106/total Rab12 ratio (n=4 experiments). C, quantification of normalized LRRK2-pSer935/total LRRK2 ratio (n=3-4 experiments). Data was analyzed by one-way ANOVA with Holm-Sidak’s multiple comparisons test. Data presented as means ± SEM; **p<0.01; ****p<0.0001 vs. LRRK2 WT.

Transient overexpression in HEK293 cells is likely to increase the Vmax of the individual LRRK2 variants compared to the endogenous LRRK2 WT enzyme. Plotting the relative Rab10-pThr73 levels vs. total LRRK2 levels suggests that Rab10 phosphorylation correlates with total LRRK2 expression levels (Fig 10A). Indeed, there was a statistically positive correlation between the two variables, (Pearson correlation coefficient; r = 0.6272, p < 0.01, with a R^2^ = 0.3934; 95% [0.2267 to 0.8463]). This was not observed for Rab12 phosphorylation (Fig 10B) and no statistically positive correlation was observed between Rab12-pSer106 levels vs. total LRRK2 levels (Pearson correlation coefficient; r = 0.3198, p > 0.05, with a R^2^ = 0.1023; 95% [-0.1729 to 0.6845]). Also, there was no significant correlation between total Rab10 or Rab12 levels and the phosphorylation at LRRK2-Ser935 (Fig 10C-D) suggesting that Rab10 or Rab12 levels do not impact LRRK2-Ser935 phosphorylation or vice versa.

**Figure 10.**
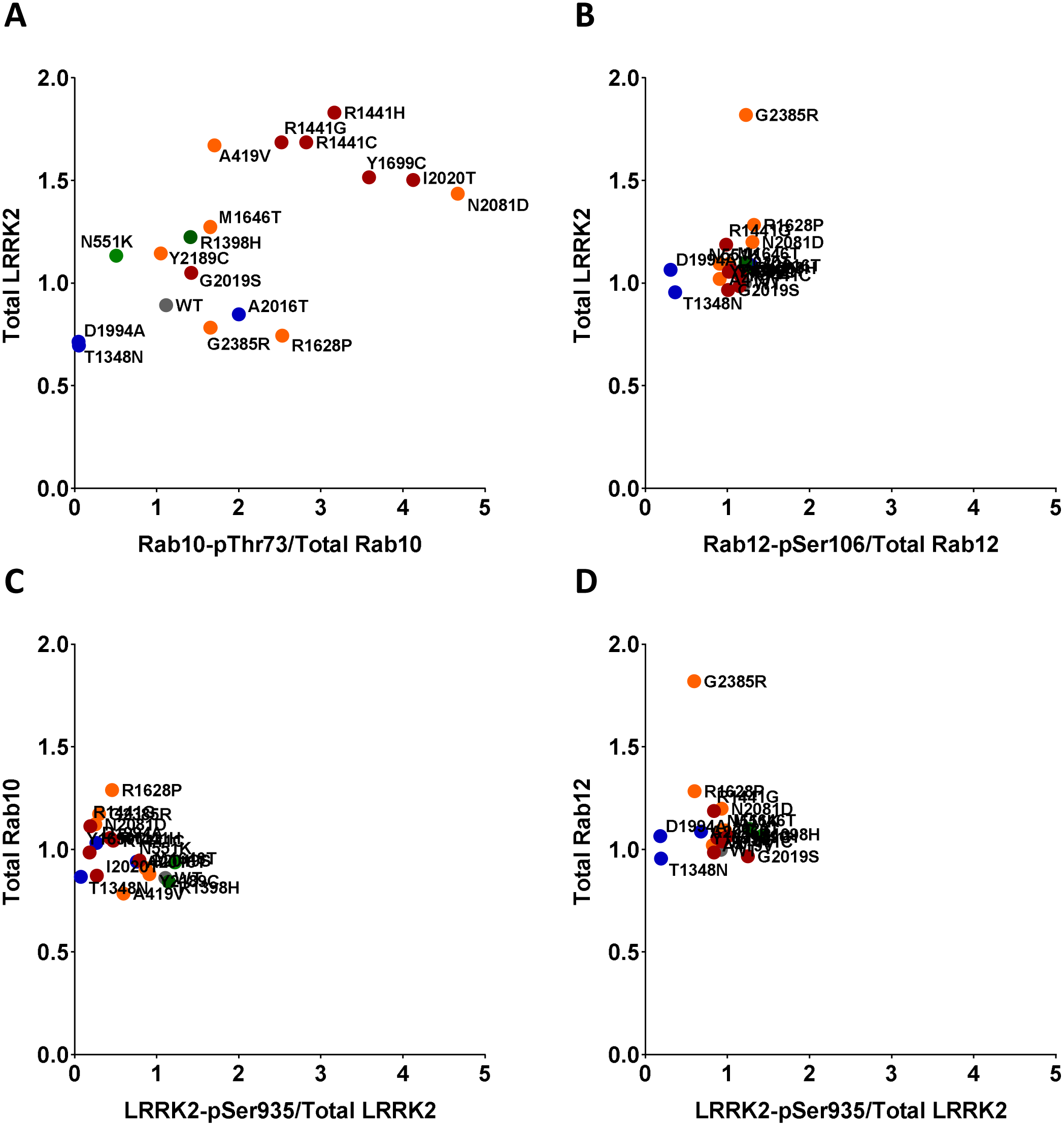
Rab10-Thr73 phosphorylation is significantly correlated with total LRRK2 expression in HEk293 cells. A, Total LRRK2 expression versus Rab10-pThr73/Total Rab10 levels. B, Total LRRK2 expression versus Rab12-pSer106/Total Rab12 levels. C, Total Rab10 versus LRRK2-pSer935/Total LRRK2 levels. D, Total Rab12 versus LRRK2-pSer935/Total LRRK2 levels.

## Discussion

Currently, no pharmacodynamic biomarker exists that can measure LRRK2 catalytic activity in human bio samples. In clinical trials investigating the therapeutic potential of LRRK2 kinase inhibition in Parkinson’s disease to obtain such a biomarker would be highly relevant. Not only for patient segmentation purposes but also for initial enrichment of PD subjects where the LRRK2 biology is likely to play a larger role. As an example, in the Caucasian, Asian and North-African Arab populations risk variants M1646T, A419V, G2385R and G2019S all have minor allele frequencies (MAF) that allow for such enrichment or segmentation. Also, the PD protective haplotype variant N551K/R1398K/K1423K is quite common (MAF ∼8%) in these populations (26). With the exception of G2019S it is not, at this stage, known whether individuals carrying common risk or protective variants have increased or decreased LRRK2 kinase activity, respectively, in areas highly relevant for Parkinson’s disease. For upcoming clinical trials investigating the therapeutic effect of LRRK2 kinase inhibition such information could be very relevant. This would allow for inclusion of patients carrying risk variants and/or excluding patients carrying protective variants thus increasing the likelihood of success. In addition, such segmentation is likely to decrease the heterogeneity of the studied patient population.

LRRK2 kinase activity-dependent markers are likely to be phosphorylation sites on LRRK2 substrate proteins. This report provides a comprehensive profiling of LRRK2 exonic variation with respect to indirect determined phosphorylation occupancies, autophosphorylation and substrate phosphorylation. Our results revealed an elevated phosphorylation site occupancy cluster in just one region of LRRK2 between the ANK- and LRR domains. All sites but Thr826 have previously been reported as cellular phosphorylation sites (53;54;56;67). In our study phosphorylation at the Ser1292 site could not be identified by the LC-MS/MS study. The most likely reason why quantitative information could not be gathered for Ser1292 is that this amino acid is in close proximity to two tryptic cleavage sites. It is therefore likely that in case of an efficient tryptic digestion the peptide comprising Ser1292 cannot be identified as the generated peptide is just too short for detection by MS. Other reasons why no phosphorylation occupancy could be determined for this position might be reasoned in poor ionization properties of eventually occurring misscleaved tryptic peptides. We identified the SFPNEMGK and IWDLPLDELHLNFDFK peptides but not LS^1292^K-containing peptides; the LRRK2 sequence around the S1292 site is as follows: SFPNEMGKLS^1292^KIWDLPLDELHLNFDFK.

No obvious disease correlation could be observed for the phosphorylation at Ser910, Ser935, Ser955 and Ser973 either using MS, MRM assays or Western Blot procedures. Our data results confirm previous reports (40). On the contrary, increased phosphorylation status at Ser1292 was observed for all the causal LRRK2 exonic variants thus confirming previous literature reports on single exonic variants (14;15). In our hands the level of phosphorylation at Ser1292 is very low, even for heterologous expression systems. The detection limit on Western Blotting is approximately 1ng for purified recombinant G2019S and above 30ng for purified recombinant LRRK2 WT (Supplementary Fig 4). For comparison, we have estimated that endogenous LRRK2 expression in HEK293 cells is ∼50pg in a 20µg homogenate (data not shown) suggesting that at least 12mg protein lysate loaded on a SDS gel would be required to detect the endogenous level of phosphorylated Ser1292 using this antibody. It is therefore highly unlikely that Western Blot based detection methods can be used to evaluate endogenous Ser1292 phosphorylation status in human bio samples from Parkinson’s diease patients.

Instead, LRRK2 phosphorylation at Rab10 might provide a better alternative. Our data suggests that LRRK2 kinase-activity dependent Rab10-Thr73 phosphorylation level follow the disease risk associated with the individual LRRK2 exonic variant. However, since some variation is observed in the overexpression system after correction for multiple comparisons the Rab10 phosphorylation at Thr73 is only statistically significant for the N2081D, I2020T and Y1699C variants. Therefore, additional experiments are warranted to substantiate the observed nominal effects. An obvious caveat is that our studies have been performed in heterologous expression systems and not on cells expressing physiological relevant LRRK2 levels. However, in order to be able to detect significant increases in Rab10 phosphorylation levels of common LRRK2 risk variants more senstitive tools or platforms have to be developed. One solution might be the development of high sensitivity MSD or Simoa assays.

Another interesting observation is the fact that N551K is less whereas R1398H is more phosphorylated on Rab10-Thr73 when compared to LRRK2 WT. These two naturally occurring SNPs are in strong linkage disequilibrium in the human genome and therefore always co-segregate thus forming a common LRRK2 haplotype that has been shown to be associated with decreased risk of Parkinson’s disease (26). Our data suggests that if LRRK2 kinase activity is the driver of disease risk the protective effect might originate from the N551K polymorphism in the ARM-ANK domains rather than as suggested by literature reports from R1398H polymorphism in the ROC-Cor domain (68). This is not observed for Rab12-pSer106. It is also dependent on LRRK2 kinase activity but, in contrast to Rab10-pThr73, it does not seem to follow the disease risk associated with the variants. It is likely, though, that Rab10 interferes with LRRK2 structure or function in a way that is different from the interaction with Rab12. In support, we observed that total levels of LRRK2 correlated well with Rab10 phosphorylation at Thr73. This was not observed for Rab12 phosphorylation at Ser106. However, it is still unknown whether the Rab10-pThr73/total LRRK2 correlation just stems from an increased LRRK2 Vmax, a changed interaction between the proteins or maybe a changed subcellular localization thus impacting both LRRK2 kinase activity dependent Rab10 phosphorylation as well as indirect LRRK2-Ser935 phosphorylation. It is also possible that the LRRK2 interaction with Rab10 increases the LRRK2 protein half-life in a way that is more pronounced for variants associated with increased risk of PD and in a way, that is not observed for Rab12. Intriguingly, the protective variant N551K shows diminished Rab10-Thr73 phosphorylation, yet, the variant gives higher total levels of protein than observed for LRRK2 WT. Evidently, LRRK2 expression levels do not give the full picture. Another possibility is that the lysine in the N551K variant is ubiquitinated thus mediating an effect on LRRK2 activity or turnover and a previous report has shown that LRRK2 dephosphorylation in HEK293 cells mediates polyubiquitination (69). Future experiments are needed to substantiate these observations.

Collectively, the data obtained for the LRRK2 exonic variants suggest that increased LRRK2 enzymatic activity correlate with increased risk of Parkinson’s disease not only in carriers of rare but also common LRRK2 exonic variants. By analogy it hereby follows that a general lowering of the LRRK2 enzymatic activity using LRRK2 kinase inhibition will decrease the risk and/or the progression of Parkinson’s disease. An important scope of future studies should be to develop more sensitive LRRK2 kinase activity detection tools to identify which patients that are eligible for clinical studies and to address how much LRRK2 kinase inhibition is required to achieve a therapeutic effect in sporadic PD patients.

## Material and Methods

### Ethical statement

All animal experiments were carried out at Genscript (China). They were also in accordance with the European Communities Council Directive (86/609/EEC) for the care and use of laboratory animals and the Danish legislation regulating animal experiments. GenScript received OLAW’s Animal Welfare Assurance on November 9, 2010. Current assurance is effective until November 20, 2020. OLAW’s Animal Welfare Assurance accentuates the responsibilities and procedures of GenScript regarding the care and use of laboratory animals. GenScript has earned AAALAC accreditation, demonstrating our commitment to responsible animal care and use through ongoing voluntary participation in AAALAC programs.

### Animals

New Zealand male rabbits at weight 2.0-2.5kg were used for the production of rabbit polyclonal antibodies at Genscript facilities (China). Rabbits were single housed under standard housing and care conditions. For immunization of rabbits, immunogens were diluted with saline or PBS and mixed with Freund’s adjuvant until emulsified and subsequently injected subcutanenously (s.c.) into 4-6 sites along the spine using a 21G hypodermic needle. For the rabbit bleeding, the skin of the rabbit ear was disinfected with alcohol and a 23”-21” syringe was used to collect blood. Terminal bleeds were performed via the carotid artery with animals under deep anesthesia. The anesthesia was induced by intramuscular injection of 40mg/kg pentobarbital. Protocols for immunization, ear margin vein bleed and carotid artery exsanguination will only expose the animals to the lowest category of pain or distress (Minimal, transient or no pain or distress). In case of sickness animals were euthanized by giving an intravenous (i.v.) injection of 100mg/kg pentobarbital through the ear vein catheter.

### Protein digestion – pilot study

An aliquot of 1 μg of flag-tagged LRRK2 G2019S mutant (Life Technologies) diluted in lysis buffer (20 mM Hepes pH 7.5, 150 mM NaCl, 0.25 % Trion X-100, 1 mM EDTA, 1 mM EGTA, containing protease- and phosphatase inhibitors). The sample was then reduced with 10 mM dithiothreitol for 30 min and subsequently alkylated in the presence of 55 mM iodoacetamide for 30 min in the dark. Thereafter, endoproteinase Lys-C (Wako) was added at an enzyme-to-substrate ratio of 1:10 and incubated for 2 h at room temperature. Samples were thereafter diluted 1:4 with 20 mM Tris-HCl pH 8.2 before adding trypsin (Promega) at an enzyme-to-substrate ratio of 1:10 followed by overnight incubation. The resulting peptide mixtures were acidified by the addition of TFA to a final concentration of 0.5 % and subsequently desalted using in-house build reversed-phase C18 STAGE-tips as described (70). For the In-Gel digest 1 μg of flag-tagged LRRK2 G2019S mutant was initially supplemented with 1x LDS buffer (Invitrogen) subsequently reduced and A distinct protein band was excised and subsequently digested with trypsin overnight (Shevchenko et al., 2006). Tryptic peptides were extracted and desalted with in-house build reversed-phase C18 STAGE-tips (70).

### Mass spectrometric analysis – pilot study

Desalted peptides were reconstituted in 0.5% formic acid and loaded onto a reverse phase analytical column (packed in-house with C18 beads), resolved by an acetonitrile gradient using an Agilent HPLC system and directly electrosprayed via a nanoelectrospray ion source into an into a LTQ-Orbitrap Velos (Thermo Fisher Scientific). The mass spectrometer was operated in the data dependent mode to automatically switch between MS full scans at a resolution R = 60,000 (at m/z = 200) with a target value of 1,000,000 counts (max. Injection time = 500 ms) and MS/MS fragmentation scans. The fifteen most intense peptide ions were selected for collision induced fragmentation in the LTQ acquired at a target value of 5000 ion counts. The resulting fragmentation spectra were recorded in the linear ion trap.

### Data processing – pilot study

All raw files were processed with the MaxQuant software suite (version 1.3.9.21) for peptide and protein identification using the human SwissProt database (version 03 2013) in which the LRRK2 sequence (Uniprot ID: Q5S007) was replaced by the LRRK2 sequences provided by life technologies (Supplementary Table 1)(71;72). The maximum mass deviations allowed for MS and MS2 peaks were 6 ppm and 0.5 Da, respectively. Carbamidomethylation of cysteine was set as a fixed modification and the oxidation of methionine and N-terminal acetylation were set as variable modifications. In a second data-processing we allowed also phosphorylation of serine, threonine and tyrosine as variable modifications. All peptides were required to have a minimum peptide length of seven amino acids and a maximum of two missed cleavages and three labeled amino acids were allowed. The false discovery rate (FDR) for protein and peptide identifications of set to 1 %.

### Data analysis – pilot study

For both protocols (In-Gel and In-Solution) the number of identified LRRK2 peptides and the corresponding sequence coverage were compared (Table 1). In addition, all LRRK2-derived phosphorylation sites that could be localized with high confidence (so called class-I phosphorylation sites (58), indicated by a localization probability ≥ 0.75 and a score difference ≥ 5 of class-I) are listed in Table 2.

### Protein digestion – LC-MS/MS phosphoproteomic and proteomic studies

For in-gel protein digest a 10 µg aliquot of each flag-tagged LRRK2 variant was employed as described above. Tryptic peptides were extracted and desalted with SepPak columns (100 mg bed volume, Waters). The resulting peptide mixtures were then aliquoted. From each LRRK2 variant an aliquot corresponding to 25 percent of the initial total protein (theoretically 2.5 μg protein/peptides) was utilized for phosphopeptide enrichment upon combining three or four different LRRK2 variants.

### Phosphopeptide enrichment - LC-MS/MS phosphoproteomic and proteomic studies

Phosphopeptides were enriched using Titanium dioxide (TiO_2_) microspheres. Briefly, Peptide samples were reconstituted in loading buffer (80% ACN 0.1% TFA containing 0.3 g/ml glycolic acid) and loaded onto previously equilibrated TiO_2_ microspheres, incubated for 10 min while shaking and loaded on C_8_ STAGEtip columns. Upon washing with loading buffer, wash buffer-I (80% ACN 1% TFA) and wash buffer-II (80% ACN 0.1% TFA) the bound phosphopeptides were eluted with 5% solution of ammonium hydroxide and dried in an Eppendorf concentrator. For MS-analysis samples were reconstituted in 0.1% formic acid.

### Peptide dephosphorylation and peptide labeling – LC-MS/MS phosphoproteomic and proteomic studies

Of each LRRK2 variant an aliquot corresponding to 5 µg of desalted tryptic peptides was dissolved in phosphatase reaction buffer (10 mM Tris-HCl (pH 8.0 at 37°C), 5 mM MgCl2, 100 mM KCl, 0.02% TritonX-100, and 0.1 mg/mL BSA) and split into two parts. The control-treated sample was supplemented with 0.5 M EDTA solution to a final concentration of 50 mM EDTA and incubated for 5 min at room temperature (25°C) and 1000 rpm using an ThermoMixer (Eppendorf). Alkaline phosphatase (FastAP, Thermo Fisher Scientific) was then added to the other sample and both samples were incubated for 1 h at 37°C and 1000 rpm. Subsequently 0.5 M EDTA solution was added to the phosphatase treated sample and both samples were incubated for another 5 min at room temperature and 1000 rpm. Note, 50 mM EDTA inactivates the phosphatase activity of the FastAP. In addition, to inactivate the thermosensitive alkaline phosphatase, samples were incubated for 15 min at 75°C. Samples were subsequently snap frozen in liquid N2 and lyophilized.

To allow for quantitative analysis the dephosphorylated and control-treated peptide samples were differentially labeled with mTRAQ reagents (light (Δ0) or heavy (Δ8); ABSciex) according to the manufacturers instructions, with the only change that a 20-fold surplus of mTRAQ reagent was used to ensure complete labeling.

The mTRAQ-labeled samples of each experiment were combined and frozen in liquid N_2_ before lyophilization. The lyophilized samples were subsequently reconstituted in 0.1% trifluoroacetic acid, desalted with in-house build reversed-phase C_18_ STAGE-tips essentially and finally reconstituted in 0.1% formic acid before MS-analysis.

### Mass spectrometric analysis – LC-MS/MS phosphoproteomic and proteomic studies

The LC-MS/MS analyses for the initial test experiments and the generation of the phosphopeptide reference dataset were performed on a Q-Exactive mass spectrometer (Thermo Fisher Scientific) while the LC-MS/MS analyses for the later experiments to determine phosphorylation occupancies were performed on a LTQ-Velos mass spectrometer (Thermo Fisher Scientific). Both mass spectrometers were equipped with an Easy-1000 UPLC (Thermo Fischer Scientific) and samples were loaded with an auto sampler onto a 40 cm fused silica emitter (New Objective) packed in-house with reversed phase material (Reprusil-Pur C18-AQ, 1.9 μm, Dr. Maisch GmbH). The bound peptides were eluted over 125 min runs and sprayed directly into the mass spectrometer using a nanoelectrospray ion source (ProxeonBiosystems). The Q-Exactive was operated in the data dependent mode to automatically switch between MS full scans at a resolution R = 70,000 (at m/z = 200) with a target value of 3,000,000 counts (max. Injection time = 45 ms). Dynamic exclusion was set to 30s and the ten most intense ions were fragmented in the octopole collision cell utilizing higher-energy collisional dissociation (HCD). Subsequent MS2 scans with a target value of 200,000 ions were collected with a maximum fill time of 120 ms and R = 35,000. The LTQ-Velos was operated in the data dependent mode to automatically switch between MS full scans at a resolution R = 60,000 (at m/z = 400) with a target value of 1,000,000 counts (max. Injection time = 500 ms) and MS/MS fragmentation scans. The fifteen most intense peptide ions were selected for collision induced fragmentation in the LTQ acquired at a target value of 5000 ion counts. The resulting fragmentation spectra were recorded in the linear ion trap.

### Data processing – LC-MS/MS phosphoproteomic and proteomic studies

All raw files were processed with the MaxQuant software suite (version 1.4.3.2) for peptide and protein identification using the human SwissProt database (version 09 2013) in which the LRRK2 sequence (UniProtKB entry: Q5S007) was replaced by the 19 DYKDDDDK-LRRK2 constructs carrying the respective amino acid variants provided by life technologies. Sequence information for all LRRK2 variants is listed in Supplementary Table 3. To generate the phosphopeptide reference dataset the raw files from a pilot study were also included.

The maximum mass deviations allowed for MS and MS2 peaks were 6 ppm and 0.5 Da for data acquired with the LTQ-Velos and 4.5 and 20 ppm for Q Exactive data. Carbamidomethylation of cysteines was set as a fixed modification and the oxidation of methionines, N-terminal acetylation and phosphorylation of serine, threonine and tyrosine were allowed as variable modifications. For all quantitative experiments the appropriate mTRAQ labels were selected as modifications. Full tryptic specificity was required, up to two missed cleavages were allowed and the minimum peptide length was set to 7 amino acids. The false discovery rate (FDR) for protein and peptide identifications of set to 1%.

### Data analysis – LC-MS/MS phosphoproteomic and proteomic studies

The list of identified phosphorylation sites was filtered for LRRK2-derived phosphorylation sites that could be localized with high confidence (so called class-I phosphorylation sites, indicated by a localization probability ≥ 0.75 and a score difference ≥ 5). Together, 43 LRRK2 derived class-I phosphosites were identified which are listed in Table 2. This dataset served as phosphopeptide reference dataset for the subsequent determination of phosphorylation site occupancies.

To calculate the phosphorylation site occupancy the following procedure was applied. For each experiment the quantified ratios for LRRK2 peptides were normalized to the median ratio of all peptides. Where appropriate these ratios were converted to match to the treatment condition where the light mTRAQ label is dephosphorylated. Ratios were log-transformed, and the mean with the corresponding 95% confidence intervals for each peptide was calculated based on a truncated t-distribution (as the stoichiometry cannot be below 0). The mean ratio was then used to calculate the percentage of phosphorylation for all LRRK2 derived peptides containing at least one serine, threonine or tyrosine residue according to the following equation: phosphorylation stoichiometry = (1- H/L) × 100% (73). Generally, the peptide with the lowest number of missed tryptic cleavage sites was used.

Initially, we visualized this data by heat map plotting of the calculated phosphorylation stoichiometry of all peptides to gain a global overview (Fig 4). Next, we performed one-way ANOVA analyses to identify phosphorylation sites with differential phosphorylation site stoichiometries among the different LRRK2 variants. This analysis was performed for all LRRK2 peptides that containing at least one serine, threonine or tyrosine residue. Site-specific information was thereupon added by matching the peptide information to the phosphopeptide reference data. ANOVA was applied to all peptides that were quantified in at least two out of four replicate experiments. The resulting p-values were corrected for multiple hypotheses testing by applying Benjamini-Hochberg’s false discovery rate (FDR). The peptides below an FDR of 5% were subjected to 2 different post-hoc tests in order to identify the variants that show significant differences in phosphorylation stoichiometry. The first, Tukey’s procedure, is a simple t-test and identifies significantly different pairs of LRRK2 variants (p<0.05). The second test compares each variant to all others in order to identify variants that behave differently to the general trend (p<0.05).

### Synthetic Peptides – MRM studies

Synthetic peptides were purchased from Thermo Fisher Scientific as crude unlabeled (PEPotec™ SRM peptide library) and heavy-labeled (Arg10/Lys8), purified (<95%, except peptide #2 73% purity) versions.

### Assay characterization – MRM studies

For assay characterization, peptides were spiked into a matrix of 0.1 μg/μl BSA, which mimicked a background of purified LRRK2 digest - as used in the main experiments. Heavy-labled peptides were spiked into a total of ten samples at various concentrations were as crude light peptides were held at a constant concentration. For all samples, 6 replicate measurements were conducted. Light to heavy transition pair ratios were used for calculation of limit of detection (LOD), coefficient of variation (CV%), and relative error (RE%). For all peptides except for the non-oxidized Ser1292 peptide, assay characterization was successful and LODs are shown in Table 2. All LOD values range in the low amol/μl concentration range, except for the peptide containing Ser935. The assay was shown to be linear above LODs in the tested concentration range. Supplementary Fig 5 and 6 show the calibration curve plot, the plot of coefficient of variation and relative residuals of the linear curve fit for the oxidized peptide M(ox)GKLSKIWDLPLDE containing Ser1292. Plots for all peptides are part of the Supplementary Data. The non-oxidized peptide containing Ser1292 proved to be too instable for characterization and furthermore caused confounding signals for characterization of the oxidized Ser1292 peptide. It also did not give a measurable signal at the highest concentration of 100 fmol/μl in the calibration curve samples as it seems to mostly oxidize. Hence, quantification for phosphorylation at the Ser1292 was done by using only the oxidized form.

### LRRK2 protein digestion - MRM studies

For the in-gel digest, 2x 10 μg of each flag-tagged LRRK2 variant was supplemented with 4x LDS buffer (Invitrogen) to a final concentration of 1x LDS. One aliquot of 10 μg of each variant was reduced and alkylated, the LRRK2 protein band of about 288 kDa was excised and digested overnight with trypsin (Promega). For a second aliquot of 10 μg of each variant, the reduction and alkylation step was omitted and samples were subjected to overnight digest with endoproteinase Glu-C. Resulting peptide mixtures were extracted and desalted with SepPak columns (100 mg bed volume, Waters).

### Phosphatase treatment - MRM studies

Desalted tryptic and GluC peptides of each LRRK2 variant were resuspended in phosphatase reaction buffer (10 mM Tris-HCl (pH 8.0 at 37°C), 5 mM MgCl2, 100 mM KCl, 0.02% TritonX-100, and 0.1 mg/mL BSA), which contained the heavy-labeled peptides to yield a concentration of 10-105 fmol/μl in the final samples. Resuspended samples were split into twelve parts each. Thus, for both digestion protocols, six phosphatase (PHOSPHO) and six control (CTRL) treatments could be performed for each of the LRRK2 variants.

The control samples were supplemented with 0.5 M EDTA solution to a final concentration of 50 mM EDTA, while samples for phosphatase treatment were supplemented with an equal volume of reaction buffer. Samples were incubated for 5 min at room temperature (25°C) and 1000 rpm using a ThermoMixer (Eppendorf). Alkaline phosphatase (FastAP, Thermo Fisher Scientific) was then added to all samples. After incubation for 1 h at 37°C and 1000 rpm, 0.5 M EDTA solution was added to all phosphatase-treated samples, while control samples were supplemented with an equal volume of reaction buffer. All samples were incubated for another 5 min at room temperature and 1000 rpm. Note that 50 mM EDTA inactivates the phosphatase activity of the FastAP. In addition, to inactivate the thermosensitive alkaline phosphatase, samples were finally incubated for 15 min at 75°C, 1000 rpm. Samples were subsequently snap-frozen in liquid N2 and lyophilized overnight. The lyophilized samples were subsequently reconstituted in 0.1 % trifluoroacetic acid, desalted with in-house build reversed-phase C18 STAGE-tips essentially as described (Rappsilber et al., 2007). High amounts of triton X-100 and glycine in the enzyme buffer made a second desalting procedure necessary. Samples were finally reconstituted in 0.1% formic acid before MS-analysis.

### Mass spectrometric analysis - MRM studies

Samples were loaded onto a reverse phase analytical column (packed in-house with C18 beads), resolved by an acetonitrile gradient using a nano-HPLC system (EasynLC, Thermo Fisher Scientific) and directly electrosprayed via a nanoelectrospray ion source into a TSQ Vantage triple quadrupole mass spectrometer (Thermo Fisher Scientific). The MS and chromatography functions were controlled by the XCalibur software. The MRM instrument method consisted of one MRM scan event over the gradient time using optimized collision energy for all peptide transitions. All data was evaluated using the Skyline software tool (version 2.6) (74).

### Data evaluation - MRM studies

For all replicate analyses of the calibration curve, the concentration of every peptide was calculated by a linear model adapted from Chang et al., 2012 (75) In order to determine the limit of detection (LOD) we apply a method initially proposed in Addona et al., 2009 (76). This approach calculates the LOD as:

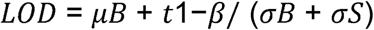

Where μB is the average calculated log-concentration of the blank, t1–β is the (1–β) percentile of the standard t distribution on f=nReplicates-1 degrees of freedom, σB is the standard deviation of calculated log-concentration of the blank and σS is the standard deviation of the calculated log-concentration at lowest actual analyte concentration. Since σB and σS can be significantly different, they are combined. As a measure of assay precision, CV% was calculated from the mean and standard deviation of the calculated concentration values for all six replicates. For calculation of endogenous peptide amounts in the samples, total area ratios between endogenous and isotopically labeled peptides were multiplied with known spike-in concentration of the labeled peptides. Phosphorylation ratios and the respective confidence intervals were estimated by bootstrapping. For each iteration, the two groups (CTRL and PHOSPHO) were randomly re-sampled with replacement and the ratio of average CTRL and average PHOSPHO was calculated. This was repeated 10.000 times. The reported ratio and upper/lower confidence bounds are represented by the median, 2.5% and 97.5% percentiles of the sampled population.

### Production and purification of a rabbit polyclonal LRRK2-pThr826 antibody

A phospho-specific antibody targeting phosphorylated LRRK2 at Thr826 was generated at Genscript using a customized “Phospho-specific Polyclonal Antibody Package”. Four rabbits were immunized at least three times using a LRRK2 phosphopeptide coupled to a KLH conjugate. The sequence used for immunization were PDK(pT)SNLRKQTNIAC. Subsequent to the 3rd immunization titer determination of phosphopeptide and nonphosphopeptide affinities guided the selection of rabbits for small-scale bleed, affinity purification as well as indirect ELISA determination of antibody specificity and selectivity. Coating antigens used for the indirect ELISA were the phosphopeptide or non-phosphopeptide. The antigens were diluted in Phosphate Buffered Saline (PBS) at pH 7.4 and used in a coating concentration and volume of 4μg/ml and 100 μl/well. The secondary antibody used for quantification was a peroxidase-conjugated goat anti-rabbit IgG. Finally, rabbits proceeded towards final boost, bleed and affinity purification of the antibodies. The ELISA results for the affinity purified LRRK2-pThr826 antibodies are listed in Supplementary Table 3. The antibody showed high specificity; however, some affinity for the non-phosphopeptide was also observed.

#### Cell culture and transfections

Plasmids encoding various LRRK2 variants as well as HA-tagged Rab10 and Rab12 protein were transiently expressed in HEK293 cells (a kind gift from Dr. JM Mathiesen, Department of Drug Design and Pharmacology, University of Copenhagen), originally obtained from ATCC (Boras, Sweden)(77), using Lipofectamine®2000 according to the manufacturer’s instructions (Thermo Fisher Scientific, US). Prior to transfections, HEK293 cells were plated at a density of 0.5×10^6^ cells/well in 12-well plates pre-coated for 1 hr at 37°C with poly-L-lysin. Subsequent to the transient transfection HEK293 cells were harvested using 1 mL cold Phosphate Buffered Saline (PBS) buffer (Invitrogen, California, US) and subsequently centrifuged at 800 g for 2 minutes. Cell pellets were resuspended and solubilized in 100 µL lysis buffer (50 mM Tris hydrochloride, 1 mM magnesium chloride, 1% Triton, 0.1% sodium dodecyl sulfate (SDS), pH 8.0) on ice for 20 minutes and then centrifuged at 20000 g for 30 minutes at 4 °C.

#### SDS-PAGE and Western Blotting of mammalian purified LRRK2 proteins

Purified LRRK2 protein was size mobility separated by SDS-polyacrylamide gel electrophoresis (SDS-PAGE) using a 3-8% Tris-Acetate gel (NuPAGE^®^ Tris-Acetate Mini Gels, Life Technologies, Paisley, UK). An amount of 10ng protein was loaded in each well for studies on Ser910, Ser935, Ser955 and Ser973 whereas in studies assessing Ser1292 phosphorylation 90ng of protein was loaded. Proteins were transferred to immobilon-FL PVDF membranes (Millipore, Billerica, US). Membranes were incubated with primary antibodies overnight at 4°C: mouse monoclonal [N241A/34] anti-LRRK2 antibody (1:2.000; NeuroMab, California, US), rabbit monoclonal [UDD1 15(3)] (ab133449) anti-pS910-LRRK2 antibody (1:1.000; RabMAb^®^, Abcam, Cambridge, UK), rabbit monoclonal [UDD2 10(12)] anti-pS935-LRRK2 antibody (1:1.000; RabMAb^®^, Abcam, Cambridge, UK), rabbit monoclonal [MJF-R11 (75-1)] (ab169521) anti-pS955-LRRK2 antibody (1:1.000; RabMAb^®^, Abcam, Cambridge, UK), rabbit monoclonal [MJF-R12 (37-1)] (ab181364) anti-pS973-LRRK2 antibody (1:1.000; RabMAb^®^, Abcam, Cambridge, UK), rabbit monoclonal [MJFR-19-7-8] (ab203181) anti-pS1292-LRRK2 antibody (1:1.000; RabMAb^®^, Abcam, Cambridge, UK). Subsequently, the membranes were washed and incubated with secondary antibodies for 1 hour at room temperature: Anti-rabbit IgG F(c) (GOAT) antibody IRDye^®^ 800CW Conjugated (1:10.000; Rockland Immunochemicals Inc., Gilbertsville, US) and anti-mouse Alexa Fluor^®^ 680 Goat anti-mouse IgM (1:20.000; Life Technologies, Paisley, UK). The proteins were visually detected by infrared imaging using Li-Cor Odyssey CLx (LI-COR, Nebraska, US). Membranes were scanned and band intensities were quantified using the Li-Cor Odyssey software (Image Studio version 3.1.4).

#### SDS-PAGE and Western Blotting of HEK293 cell lysates

The solubilized crude cell culture lysate was size mobility separated by SDS-polyacrylamide gel electrophoresis (SDS-PAGE). LRRK2 was separated on a 3-8% Tris-Acetate gel (NuPAGE^®^ Tris-Acetate Mini Gels, Life Technologies, Paisley, UK) and Rab10 and Rab12 were separated using a NuPAGE^®^ Novex 4-12% Bis-Tris Gel (Invitrogen, California, US). An amount of 15 µg total protein was loaded in each well. The subsequent protein transfer, antibody incubations, membrane washing, visual detection and data analysis were performed as above mentioned. The following primary antibodies were used: anti-LRRK2 antibodies as mentioned above, mouse anti-HA monoclonal antibody [clone HA-7] (1:10,000; Santa Cruz Biotechnology, Texas, US), rabbit anti-Rab10-pThr73 polyclonal [#5981] (1:2,000; H. Lundbeck A/S, DK) and rabbit anti-Rab12-Ser106 polyclonal antibody [#5919] (1:5,000; H. Lundbeck A/S, DK) (66).

#### Data analysis and statistics

Data and statistical analyses were performed using Prism 5 (GraphPad Software, USA) Data were analyzed by either 1-way or 2-way analysis of variance (ANOVA) or by t-test. Post-hoc tests following ANOVAs were conducted using Dunnett’s or Holm-Sidak’s multiple comparisons test. Two-tailed levels of significance were used and p<0.05 was considered statistically significant.

## Acknowledgments

We are grateful to Kirsten Olesen and Louise Tarpø for their skilled technical assistance and to Boris Xu from Genscript for helpful assistance with information on animal protocols and procedures.

## Supporting information

**S1 Table. LRRK2 phospho-peptide mapping.**

**S2 Table. Mammalian purified LRRK2 protein batches.**

**S1 Fig. LRRK2 phosphorylation site stochiometries**. *A*, Ser910; *B*, Ser935; *C*, Ser955 and *D*, Ser973 sites.

**S2 Fig. LRRK2 phosphorylation site stochiometry for the Ser826 site.**

**S3 Fig. 14-3-3 peptides co-purified with individual LRRK2 variants.** A, YWHAE (epsilon); B, YWHAQ (theta); C, YWHAZ (zeta); D, YWHAG (gamma); E, YWHAB (beta) and F, YWAH (eta).

**S3 Table. Indirect ELISA results for LRRK2-pThr826 polyclonal antibody.**

**S4 Fig. Sensitivity of Ser1292 Western Blot assay.** A, Western Blot images from SDS-PAGE concentration curves of mammalian purified LRRK2 WT (*top panel*), D1994A (*middle panel*) and G2019S (*bottom panel*) proteins. B, quantification of the relative LRRK2-pSer1292 levels (pSer1292/total LRRK2).

**S5 Fig. Calibration curves, coefficient of variation, and relative residuals of linear fit for all LRRK2 peptides.** The peptides for the following phosphorylation sites were assessed: *A*, Ser973; *B*, Ser935 and *C*, Ser955. *Top*, calibration curve plot of actual concentration of heavy peptide against calculated concentration using the curve equation generated by linear fit; the calculated limit of detection (LOD) is indicated by the bold vertical line. *Middle,* plot of coefficient of variation of replicate samples for all concentration points. *Bottom*, relative residuals of linear calibration curve fit against concentration.

**S6 Fig. Calibration curves, coefficient of variation, and relative residuals of linear fit for all LRRK2 peptides.** The peptides for the following phosphorylation sites were assessed: *A*, Ser1292; *B*, Ser910 and *C*, Thr826. *Top*, calibration curve plot of actual concentration of heavy peptide against calculated concentration using the curve equation generated by linear fit; the calculated limit of detection (LOD) is indicated by the bold vertical line. *Middle,* plot of coefficient of variation of replicate samples for all concentration points. *Bottom*, relative residuals of linear calibration curve fit against concentration.

## References

(1) Aasly JO, Vilarino-Guell C, Dachsel JC, Webber PJ, West AB, Haugarvoll K, et al. Novel pathogenic LRRK2 p. Asn1437His substitution in familial Parkinson’s disease. Mov Disord 2010 Oct 15;25(13): 2156-63.

(2) Funayama M, Hasegawa K, Ohta E, Kawashima N, Komiyama M, Kowa H, et al. An LRRK2 mutation as a cause for the parkinsonism in the original PARK8 family. Ann Neurol 2005 Jun;57(6): 918-21.

(3) Funayama M, Hasegawa K, Kowa H, Saito M, Tsuji S, Obata F. A new locus for Parkinson’s disease (PARK8) maps to chromosome 12p11.2-q13.1. Ann Neurol 2002 Mar;51(3): 296-301.

(4) Kachergus J, Mata IF, Hulihan M, Taylor JP, Lincoln S, Aasly J, et al. Identification of a novel LRRK2 mutation linked to autosomal dominant parkinsonism: evidence of a common founder across European populations. Am J Hum Genet 2005 Apr;76(4): 672-80.

(5) Gilks WP, Abou-Sleiman PM, Gandhi S, Jain S, Singleton A, Lees AJ, et al. A common LRRK2 mutation in idiopathic Parkinson’s disease. Lancet 2005 Jan 29;365(9457): 415-6.

(6) Di FA, Rohe CF, Ferreira J, Chien HF, Vacca L, Stocchi F, et al. A frequent LRRK2 gene mutation associated with autosomal dominant Parkinson’s disease. Lancet 2005 Jan 29;365(9457): 412-5.

(7) Nichols WC, Pankratz N, Hernandez D, Paisan-Ruiz C, Jain S, Halter CA, et al. Genetic screening for a single common LRRK2 mutation in familial Parkinson’s disease. Lancet 2005 Jan 29;365(9457): 410-2.

(8) Zimprich A, Biskup S, Leitner P, Lichtner P, Farrer M, Lincoln S, et al. Mutations in LRRK2 cause autosomal-dominant parkinsonism with pleomorphic pathology. Neuron 2004 Nov 18;44(4): 601-7.

(9) Lorenzo-Betancor O, Samaranch L, Ezquerra M, Tolosa E, Lorenzo E, Irigoyen J, et al. LRRK2 haplotype-sharing analysis in Parkinson’s disease reveals a novel p.S1761R mutation. Mov Disord 2012 Jan;27(1): 146-51.

(10) Mata IF, Davis MY, Lopez AN, Dorschner MO, Martinez E, Yearout D, et al. The discovery of LRRK2 p.R1441S, a novel mutation for Parkinson’s disease, adds to the complexity of a mutational hotspot. Am J Med Genet B Neuropsychiatr Genet 2016 Apr 25.

(11) Paisan-Ruiz C, Jain S, Evans EW, Gilks WP, Simon J, van der Brug M, et al. Cloning of the gene containing mutations that cause PARK8-linked Parkinson’s disease. Neuron 2004 Nov 18;44(4): 595-600.

(12) Zabetian CP, Samii A, Mosley AD, Roberts JW, Leis BC, Yearout D, et al. A clinic-based study of the LRRK2 gene in Parkinson disease yields new mutations. Neurology 2005 Sep 13;65(5): 741-4.

(13) Jaleel M, Nichols RJ, Deak M, Campbell DG, Gillardon F, Knebel A, et al. LRRK2 phosphorylates moesin at threonine-558: characterization of how Parkinson’s disease mutants affect kinase activity. Biochem J 2007 Jul 15;405(2): 307-17.

(14) Sheng Z, Zhang S, Bustos D, Kleinheinz T, Le Pichon CE, Dominguez SL, et al. Ser1292 autophosphorylation is an indicator of LRRK2 kinase activity and contributes to the cellular effects of PD mutations. Sci Transl Med 2012 Dec 12;4(164): 164ra161.

(15) Henry AG, Aghamohammadzadeh S, Samaroo H, Chen Y, Mou K, Needle E, et al. Pathogenic LRRK2 mutations, through increased kinase activity, produce enlarged lysosomes with reduced degradative capacity and increase ATP13A2 expression. Hum Mol Genet 2015 Nov 1;24(21): 6013-28.

(16) Mata IF, Checkoway H, Hutter CM, Samii A, Roberts JW, Kim HM, et al. Common variation in the LRRK2 gene is a risk factor for Parkinson’s disease. Mov Disord 2012 Dec;27(14): 1822-5.

(17) Satake W, Nakabayashi Y, Mizuta I, Hirota Y, Ito C, Kubo M, et al. Genome-wide association study identifies common variants at four loci as genetic risk factors for Parkinson’s disease. Nat Genet 2009 Dec;41(12): 1303-7.

(18) Ross OA, Farrer MJ, Wu RM. Common variants in Parkinson’s disease. Mov Disord 2007 Apr 30;22(6): 899-900.

(19) Skipper L, Li Y, Bonnard C, Pavanni R, Yih Y, Chua E, et al. Comprehensive evaluation of common genetic variation within LRRK2 reveals evidence for association with sporadic Parkinson’s disease. Hum Mol Genet 2005 Dec 1;14(23): 3549-56.

(20) Funayama M, Li Y, Tomiyama H, Yoshino H, Imamichi Y, Yamamoto M, et al. Leucine-rich repeat kinase 2 G2385R variant is a risk factor for Parkinson disease in Asian population. Neuroreport 2007 Feb 12;18(3): 273-5.

(21) Farrer MJ, Stone JT, Lin CH, Dachsel JC, Hulihan MM, Haugarvoll K, et al. Lrrk2 G2385R is an ancestral risk factor for Parkinson’s disease in Asia. Parkinsonism Relat Disord 2007 Mar;13(2): 89-92.

(22) Tan EK, Tan LC, Lim HQ, Li R, Tang M, Yih Y, et al. LRRK2 R1628P increases risk of Parkinson’s disease: replication evidence. Hum Genet 2008 Oct;124(3): 287-8.

(23) Ross OA, Wu YR, Lee MC, Funayama M, Chen ML, Soto AI, et al. Analysis of Lrrk2 R1628P as a risk factor for Parkinson’s disease. Ann Neurol 2008 Jul;64(1): 88-92.

(24) Li K, Tang BS, Liu ZH, Kang JF, Zhang Y, Shen L, et al. LRRK2 A419V variant is a risk factor for Parkinson’s disease in Asian population. Neurobiol Aging 2015 Oct;36(10): 2908-5.

(25) Li NN, Tan EK, Chang XL, Mao XY, Zhang JH, Zhao DM, et al. Genetic analysis of LRRK2 A419V variant in ethnic Chinese. Neurobiol Aging 2012 Aug;33(8): 1849-3.

(26) Ross OA, Soto-Ortolaza AI, Heckman MG, Aasly JO, Abahuni N, Annesi G, et al. Association of LRRK2 exonic variants with susceptibility to Parkinson’s disease: a case-control study. Lancet Neurol 2011 Oct;10(10): 898-908.

(27) Chen L, Zhang S, Liu Y, Hong H, Wang H, Zheng Y, et al. LRRK2 R1398H polymorphism is associated with decreased risk of Parkinson’s disease in a Han Chinese population. Parkinsonism Relat Disord 2011 May;17(4): 291-2.

(28) Christensen KV, Smith GP, Williamson DS. Development of LRRK2 Inhibitors for the Treatment of Parkinson’s Disease. Prog Med Chem 2017;56:37-80.

(29) Estrada AA, Sweeney ZK. Chemical Biology of Leucine-Rich Repeat Kinase 2 (LRRK2) Inhibitors. J Med Chem 2015 Sep 10;58(17): 6733-46.

(30) Liu G, Sgobio C, Gu X, Sun L, Lin X, Yu J, et al. Selective expression of Parkinson’s disease-related Leucine-rich repeat kinase 2 G2019S missense mutation in midbrain dopaminergic neurons impairs dopamine release and dopaminergic gene expression. Hum Mol Genet 2015 Sep 15;24(18): 5299-312.

(31) Yue M, Hinkle KM, Davies P, Trushina E, Fiesel FC, Christenson TA, et al. Progressive dopaminergic alterations and mitochondrial abnormalities in LRRK2 G2019S knock-in mice. Neurobiol Dis 2015 Jun;78:172-95.

(32) Garcia-Miralles M, Coomaraswamy J, Habig K, Herzig MC, Funk N, Gillardon F, et al. No dopamine cell loss or changes in cytoskeleton function in transgenic mice expressing physiological levels of wild type or G2019S mutant LRRK2 and in human fibroblasts. PLoS One 2015;10(4):e0118947.

(33) Lee JW, Tapias V, Di MR, Greenamyre JT, Cannon JR. Behavioral, neurochemical, and pathologic alterations in bacterial artificial chromosome transgenic G2019S leucine-rich repeated kinase 2 rats. Neurobiol Aging 2015 Jan;36(1): 505-18.

(34) Walker MD, Volta M, Cataldi S, Dinelle K, Beccano-Kelly D, Munsie L, et al. Behavioral deficits and striatal DA signaling in LRRK2 p.G2019S transgenic rats: a multimodal investigation including PET neuroimaging. J Parkinsons Dis 2014;4(3): 483-98.

(35) Chou JS, Chen CY, Chen YL, Weng YH, Yeh TH, Lu CS, et al. (G2019S) LRRK2 causes early-phase dysfunction of SNpc dopaminergic neurons and impairment of corticostriatal long-term depression in the PD transgenic mouse. Neurobiol Dis 2014 Aug;68:190-9.

(36) Hindle SJ, Elliott CJ. Spread of neuronal degeneration in a dopaminergic, Lrrk-G2019S model of Parkinson disease. Autophagy 2013 Jun 1;9(6): 936-8.

(37) Chen CY, Weng YH, Chien KY, Lin KJ, Yeh TH, Cheng YP, et al. (G2019S) LRRK2 activates MKK4-JNK pathway and causes degeneration of SN dopaminergic neurons in a transgenic mouse model of PD. Cell Death Differ 2012 Oct;19(10): 1623-33.

(38) Ramonet D, Daher JP, Lin BM, Stafa K, Kim J, Banerjee R, et al. Dopaminergic neuronal loss, reduced neurite complexity and autophagic abnormalities in transgenic mice expressing G2019S mutant LRRK2. PLoS One 2011;6(4):e18568.

(39) Dusonchet J, Kochubey O, Stafa K, Young SM, Jr, Zufferey R, Moore DJ, et al. A rat model of progressive nigral neurodegeneration induced by the Parkinson’s disease-associated G2019S mutation in LRRK2. J Neurosci 2011 Jan 19;31(3): 907-12.

(40) Li X, Patel JC, Wang J, Avshalumov MV, Nicholson C, Buxbaum JD, et al. Enhanced striatal dopamine transmission and motor performance with LRRK2 overexpression in mice is eliminated by familial Parkinson’s disease mutation G2019S. J Neurosci 2010 Feb 3;30(5): 1788-97.

(41) Volpicelli-Daley LA, Abdelmotilib H, Liu Z, Stoyka L, Daher JP, Milnerwood AJ, et al. G2019SLRRK2 Expression Augments alpha-Synuclein Sequestration into Inclusions in Neurons. J Neurosci 2016 Jul 13;36(28): 7415-27.

(42) Daher JP, Abdelmotilib HA, Hu X, Volpicelli-Daley LA, Moehle MS, Fraser KB, et al. Leucine-rich Repeat Kinase 2 (LRRK2) Pharmacological Inhibition Abates alpha-Synuclein Gene-induced Neurodegeneration. J Biol Chem 2015 Aug 7;290(32): 19433-44.

(43) Daher JP, Volpicelli-Daley LA, Blackburn JP, Moehle MS, West AB. Abrogation of alpha-synuclein-mediated dopaminergic neurodegeneration in LRRK2-deficient rats. Proc Natl Acad Sci U S A 2014 Jun 24;111(25): 9289-94.

(44) Steger M, Tonelli F, Ito G, Davies P, Trost M, Vetter M, et al. Phosphoproteomics reveals that Parkinson’s disease kinase LRRK2 regulates a subset of Rab GTPases. Elife 2016;5.

(45) Zheng Y, Liu Y, Wu Q, Hong H, Zhou H, Chen J, et al. Confirmation of LRRK2 S1647T variant as a risk factor for Parkinson’s disease in southern China. Eur J Neurol 2011 Mar;18(3): 538-40.

(46) Heckman MG, Schottlaender L, Soto-Ortolaza AI, Diehl NN, Rayaprolu S, Ogaki K, et al. LRRK2 exonic variants and risk of multiple system atrophy. Neurology 2014 Dec 9;83(24): 2256-61.

(47) Refai FS, Ng SH, Tan EK. Evaluating LRRK2 genetic variants with unclear pathogenicity. Biomed Res Int 2015;2015:678701.

(48) Jasinska-Myga B, Kachergus J, Vilarino-Guell C, Wider C, Soto-Ortolaza AI, Kefi M, et al. Comprehensive sequencing of the LRRK2 gene in patients with familial Parkinson’s disease from North Africa. Mov Disord 2010 Oct 15;25(13): 2052-8.

(49) Greggio E, Jain S, Kingsbury A, Bandopadhyay R, Lewis P, Kaganovich A, et al. Kinase activity is required for the toxic effects of mutant LRRK2/dardarin. Neurobiol Dis 2006 Aug;23(2): 329-41.

(50) West AB, Moore DJ, Biskup S, Bugayenko A, Smith WW, Ross CA, et al. Parkinson’s disease-associated mutations in leucine-rich repeat kinase 2 augment kinase activity. Proc Natl Acad Sci U S A 2005 Nov 15;102(46): 16842-7.

(51) Ito G, Okai T, Fujino G, Takeda K, Ichijo H, Katada T, et al. GTP binding is essential to the protein kinase activity of LRRK2, a causative gene product for familial Parkinson’s disease. Biochemistry 2007 Feb 6;46(5): 1380-8.

(52) Nichols RJ, Dzamko N, Hutti JE, Cantley LC, Deak M, Moran J, et al. Substrate specificity and inhibitors of LRRK2, a protein kinase mutated in Parkinson’s disease. Biochem J 2009 Nov 15;424(1): 47-60.

(53) Dzamko N, Deak M, Hentati F, Reith AD, Prescott AR, Alessi DR, et al. Inhibition of LRRK2 kinase activity leads to dephosphorylation of Ser(910)/Ser(935), disruption of 14-3-3 binding and altered cytoplasmic localization. Biochem J 2010 Sep 15;430(3): 405-13.

(54) Nichols RJ, Dzamko N, Morrice NA, Campbell DG, Deak M, Ordureau A, et al. 14-3-3 binding to LRRK2 is disrupted by multiple Parkinson’s disease-associated mutations and regulates cytoplasmic localization. Biochem J 2010 Sep 15;430(3): 393-404.

(55) Reynolds A, Doggett EA, Riddle SM, Lebakken CS, Nichols RJ. LRRK2 kinase activity and biology are not uniformly predicted by its autophosphorylation and cellular phosphorylation site status. Front Mol Neurosci 2014;7:54.

(56) Zhao J, Hermanson SB, Carlson CB, Riddle SM, Vogel KW, Bi K, et al. Pharmacological inhibition of LRRK2 cellular phosphorylation sites provides insight into LRRK2 biology. Biochem Soc Trans 2012 Oct;40(5): 1158-62.

(57) Delbroek L, Van KK, Steegmans L, da CR, Mandemakers W, Daneels G, et al. Development of an enzyme-linked immunosorbent assay for detection of cellular and in vivo LRRK2 S935 phosphorylation. J Pharm Biomed Anal 2013 Mar 25;76:49-58.

(58) Olsen JV, Blagoev B, Gnad F, Macek B, Kumar C, Mortensen P, et al. Global, in vivo, and site-specific phosphorylation dynamics in signaling networks. Cell 2006 Nov 3;127(3): 635-48.

(59) Kamikawaji S, Ito G, Iwatsubo T. Identification of the autophosphorylation sites of LRRK2. Biochemistry 2009 Nov 24;48(46): 10963-75.

(60) Pungaliya PP, Bai Y, Lipinski K, Anand VS, Sen S, Brown EL, et al. Identification and characterization of a leucine-rich repeat kinase 2 (LRRK2) consensus phosphorylation motif. PLoS One 2010;5(10):e13672.

(61) Gloeckner CJ, Boldt K, von ZF, Helm S, Wiesent L, Sarioglu H, et al. Phosphopeptide analysis reveals two discrete clusters of phosphorylation in the N-terminus and the Roc domain of the Parkinson-disease associated protein kinase LRRK2. J Proteome Res 2010 Apr 5;9(4): 1738-45.

(62) Muda K, Bertinetti D, Gesellchen F, Hermann JS, von ZF, Geerlof A, et al. Parkinson-related LRRK2 mutation R1441C/G/H impairs PKA phosphorylation of LRRK2 and disrupts its interaction with 14-3-3. Proc Natl Acad Sci U S A 2014 Jan 7;111(1):E34-E43.

(63) Lobbestael E, Baekelandt V, Taymans JM. Phosphorylation of LRRK2: from kinase to substrate. Biochem Soc Trans 2012 Oct;40(5): 1102-10.

(64) Davies P, Hinkle KM, Sukar NN, Sepulveda B, Mesias R, Serrano G, et al. Comprehensive characterization and optimization of anti-LRRK2 (leucine-rich repeat kinase 2) monoclonal antibodies. Biochem J 2013 Jul 1;453(1): 101-13.

(65) Fraser KB, Moehle MS, Alcalay RN, West AB. Urinary LRRK2 phosphorylation predicts parkinsonian phenotypes in G2019S LRRK2 carriers. Neurology 2016 Mar 15;86(11): 994-9.

(66) Thirstrup K, Dachsel JC, Oppermann FS, Williamson DS, Smith GP, Fog K, et al. Selective LRRK2 kinase inhibition reduces phosphorylation of endogenous Rab10 and Rab12 in human peripheral mononuclear blood cells. Sci Rep 2017 Aug 31;7(1): 10300.

(67) Doggett EA, Zhao J, Mork CN, Hu D, Nichols RJ. Phosphorylation of LRRK2 serines 955 and 973 is disrupted by Parkinson’s disease mutations and LRRK2 pharmacological inhibition. J Neurochem 2012 Jan;120(1): 37-45.

(68) Heckman MG, Elbaz A, Soto-Ortolaza AI, Serie DJ, Aasly JO, Annesi G, et al. Protective effect of LRRK2 p.R1398H on risk of Parkinson’s disease is independent of MAPT and SNCA variants. Neurobiol Aging 2014 Jan;35(1): 266-14.

(69) Zhao J, Molitor TP, Langston JW, Nichols RJ. LRRK2 dephosphorylation increases its ubiquitination. Biochem J 2015 Jul 1;469(1): 107-20.

(70) Rappsilber J, Mann M, Ishihama Y. Protocol for micro-purification, enrichment, prefractionation and storage of peptides for proteomics using StageTips. Nat Protoc 2007;2(8): 1896-906.

(71) Cox J, Mann M. MaxQuant enables high peptide identification rates, individualized p.p.b.-range mass accuracies and proteome-wide protein quantification. Nat Biotechnol 2008 Dec;26(12): 1367-72.

(72) Olsen JV, Vermeulen M, Santamaria A, Kumar C, Miller ML, Jensen LJ, et al. Quantitative phosphoproteomics reveals widespread full phosphorylation site occupancy during mitosis. Sci Signal 2010 Jan 12;3(104):ra3.

(73) Wu R, Haas W, Dephoure N, Huttlin EL, Zhai B, Sowa ME, et al. A large-scale method to measure absolute protein phosphorylation stoichiometries. Nat Methods 2011 Jul 3;8(8): 677-83.

(74) MacLean B, Tomazela DM, Shulman N, Chambers M, Finney GL, Frewen B, et al. Skyline: an open source document editor for creating and analyzing targeted proteomics experiments. Bioinformatics 2010 Apr 1;26(7): 966-8.

(75) Chang CY, Picotti P, Huttenhain R, Heinzelmann-Schwarz V, Jovanovic M, Aebersold R, et al. Protein significance analysis in selected reaction monitoring (SRM) measurements. Mol Cell Proteomics 2012 Apr;11(4):M111.

(76) Addona TA, Abbatiello SE, Schilling B, Skates SJ, Mani DR, Bunk DM, et al. Multi-site assessment of the precision and reproducibility of multiple reaction monitoring-based measurements of proteins in plasma. Nat Biotechnol 2009 Jul;27(7): 633-41.

(77) Roed SN, Wismann P, Underwood CR, Kulahin N, Iversen H, Cappelen KA, et al. Real-time trafficking and signaling of the glucagon-like peptide-1 receptor. Mol Cell Endocrinol 2014 Feb 15;382(2): 938-49.

(78) Singleton A, Hardy J. A generalizable hypothesis for the genetic architecture of disease: pleomorphic risk loci. Hum Mol Genet 2011 Oct 15;20(R2):R158-R162.

(79) Lin CH, Wu RM, Tai CH, Chen ML, Hu FC. Lrrk2 S1647T and BDNF V66M interact with environmental factors to increase risk of Parkinson’s disease. Parkinsonism Relat Disord 2011 Feb;17(2): 84-8.

(80) Hernandez DG, Paisan-Ruiz C, McInerney-Leo A, Jain S, Meyer-Lindenberg A, Evans EW, et al. Clinical and positron emission tomography of Parkinson’s disease caused by LRRK2. Ann Neurol 2005 Mar;57(3): 453-6.

(81) Nuytemans K, Rademakers R, Theuns J, Pals P, Engelborghs S, Pickut B, et al. Founder mutation p.R1441C in the leucine-rich repeat kinase 2 gene in Belgian Parkinson’s disease patients. Eur J Hum Genet 2008 Apr;16(4): 471-9.

(82) Mata IF, Kachergus JM, Taylor JP, Lincoln S, Aasly J, Lynch T, et al. Lrrk2 pathogenic substitutions in Parkinson’s disease. Neurogenetics 2005 Dec;6(4): 171-7.

